# A temperature-sensitive metabolic valve and a transcriptional feedback loop drive rapid homeoviscous adaptation in *Escherichia coli*

**DOI:** 10.1101/2023.07.10.548422

**Authors:** Loles Hoogerland, Stefan van den Berg, Adja Zoumaro-Djayoon, Esther Geurken, Flora Yang, Frank Bruggeman, Gregory Bokinsky

**Affiliations:** Department of Bionanoscience, Kavli Institute of Nanoscience, Delft University of Technology, Delft, The Netherlands; Systems Biology Lab, AIMMS/ALIFE, Vrije Universiteit Amsterdam, Amsterdam, The Netherlands

**Author notes:** These authors contributed equally to the manuscript. Materials and correspondence.

## Abstract

All free-living microorganisms homeostatically maintain the fluidity of their membranes by adapting lipid composition to environmental temperatures. A quantitative description of how organisms maintain constant fluidity at all growth temperatures has not been achieved. By quantifying both enzymes and metabolic intermediates of the *Escherichia coli* fatty acid and phospholipid synthesis pathways, we discover how *E. coli* measures steady-state temperature and restores optimal membrane fluidity within a single generation after temperature shocks. The first element of the system is a temperature-sensitive metabolic valve that allocates flux between the saturated and unsaturated fatty acid synthesis pathways. The second element is a transcription-based negative feedback loop that counteracts the temperature-sensitive valve. The combination of these elements accelerates membrane adaptation by causing a transient overshoot in the synthesis of saturated or unsaturated fatty acids following temperature shocks. This overshoot strategy accelerates membrane adaptation, and is comparable to increasing the temperature of a water bath by adding water that is excessively hot rather than adding water at the desired temperature. These properties are captured in a quantitative model, which we further use to show how hard-wired parameters calibrate the system to generate membrane compositions that maintain constant fluidity across a wide range of temperatures. We hypothesize that core design features of the *E. coli* system will prove to be ubiquitous features of homeoviscous adaptation systems.

## Introduction

Lipid membranes provide living cells with a semi-permeable barrier, a platform for cell wall assembly, and a channel for electron transport. These and many other essential functions are strongly affected by the membrane fluidity (or viscosity), a physical property that is highly sensitive to temperature (1). Specifically, low temperatures reduce membrane fluidity by increasing the packing of membrane lipids. Organisms counteract the effects of temperature by varying the proportion of lipids that disrupt membrane packing such as unsaturated or branched-chain fatty acids **(Figure 1A)**. This response, known as homeoviscous adaptation, precisely maintains the viscosity of cell membranes at a fixed value across all growth temperatures (2). A variety of membrane sensors, metabolic pathways, and transcriptional regulators required for homeoviscous adaptation have been described in prokaryotic and eukaryotic microorganisms (3–5) and multicellular organisms (6). How these components stabilize membrane fluidity at all growth temperatures is poorly understood.

**Figure 1.**
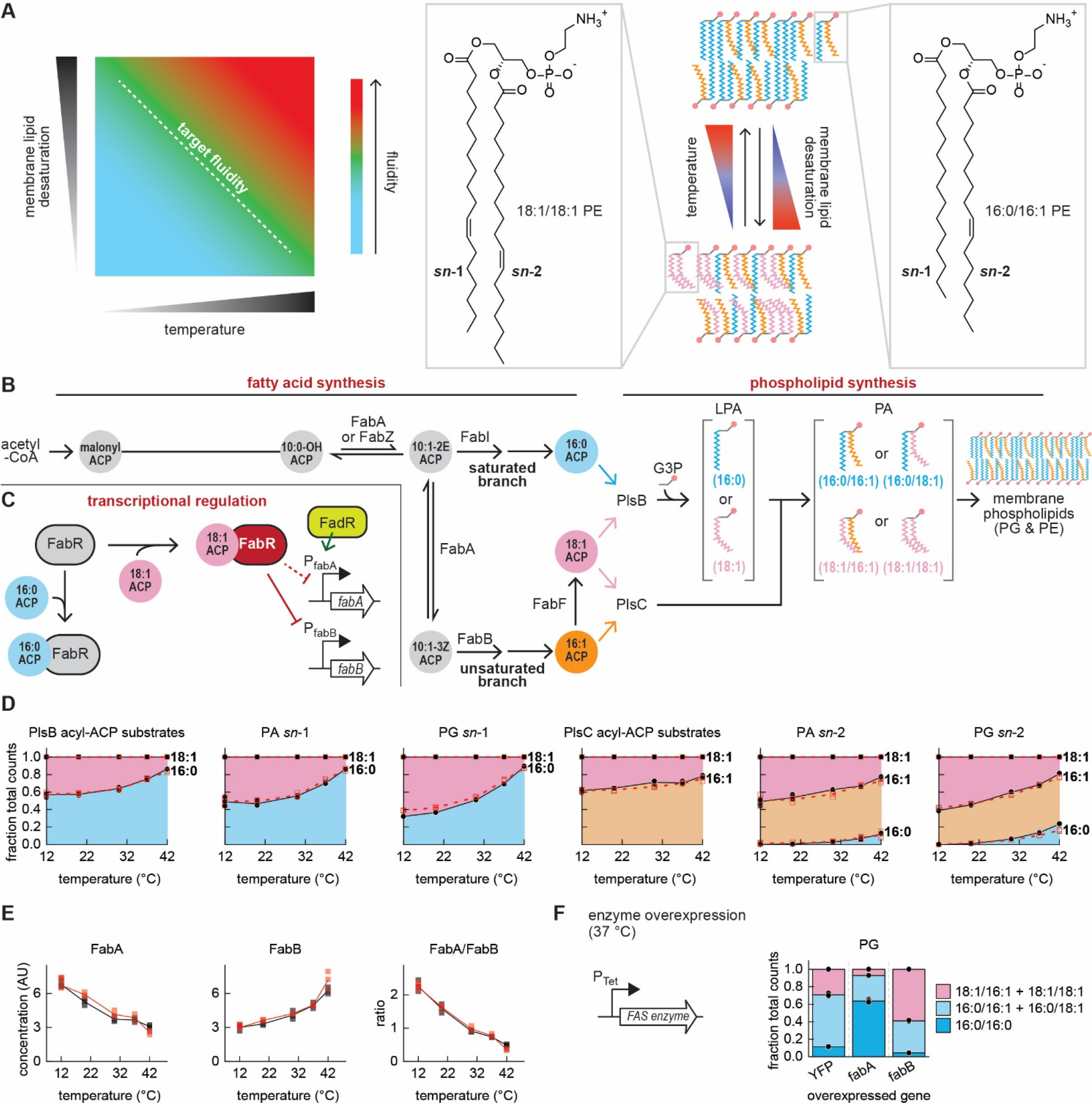
Homeoviscous adaptation and membrane metabolism in *E. coli*. **A.** *E. coli* maintains membrane fluidity by titrating the fraction of membrane phospholipids bearing two unsaturated fatty acids. **B.** The *E. coli* fatty acid synthesis pathway splits into saturated and unsaturated branches at the enoyl-acyl-ACP intermediate C10:1-2*E* ACP. Reduction by the enoyl-acyl-ACP reductase FabI initiates synthesis of saturated fatty acid thioesters (primarily C16:0 ACP). Alternatively, reversible isomerization of C10:1-2*E* ACP by the bifunctional hydroxyl-acyl-ACP dehydratase/enoyl-acyl-ACP isomerase FabA generates C10:1-3*Z* ACP, a substrate for the β-keto-acyl-ACP synthase FabB. The FabB reaction initiates synthesis of the unsaturated fatty acid thioester C16:1 ACP. A fraction of the C16:1 ACP pool is elongated by the β-keto-acyl synthase enzyme FabF and ultimately to C18:1 ACP. Phospholipids are synthesized by PlsB and PlsC, which transfer acyl groups to glycerol-3- phosphate (G3P) to produce phosphatidic acid (PA). The headgroup of PA is further modified to yield membrane phospholipids phosphatidylglycerol (PG) and phosphatidylethanolamine (PE). The complete *E. coli* fatty acid and phospholipid synthesis pathways are depicted in **Supplemental** Figure 1. **C.** The transcriptional regulator FabR in complex with C18:1 ACP reduces unsaturated fatty acid synthesis by repressing FabB. FabA expression is primarily controlled by the transcriptional regulator FadR, which adapts fatty acid synthesis in the presence of fatty acyl-CoA generated from exogenous fatty acids (31). **D.** LCMS quantification of long-chain acyl-ACP, PA, and PG. Symbols depicted are LCMS counts obtained from 3 measurements of 2 independent cultures (distinguished by solid black and open red symbols). Lines indicate the average obtained for the cultures (solid black and dashed red lines). Values are normalized to fraction of total LCMS counts per unit biomass (determined by optical density (OD)), which approximates relative concentrations. Areas represent total LCMS counts for phospholipid species bearing acyl chains at either the *sn-*1 or *sn-*2 positions as indicated. **E.** Relative abundance of FabA and FabB enzymes in cultures maintained at 5 temperatures. 3 measurements (symbols) and their average (line) are depicted for 2 independent cultures at each temperature. **F.** Phospholipid compositions of strains overexpressing YFP (control), FabA, and FabB enzymes. Bars depict average of 3 measurements (depicted by symbols) from one culture each.

In *Bacillus subtilis* and model eukaryotes, membrane composition control is directly linked with membrane fluidity. Membrane properties are monitored by integral protein sensors that interact with transcriptional pathways (7, 8). These pathways control expression of enzymes that change membrane composition by modifying existing membrane lipids (*B. subtilis* (9)) or lipid precursors (*Saccharomyces cerevisiae* (10)). In contrast, membrane fluidity and composition control are indirectly connected in *Escherichia coli. E. coli* does not monitor membrane fluidity and controls its membrane composition by varying the composition of the pool of fatty acid precursors from which phospholipids are synthesized **(Figure 1B)** (11). While transcriptional regulators that affect the composition of the fatty acid pool have been identified **(Figure 1C)** (12), temperature also directly controls the fatty acid pool composition via a transcription-independent mechanism (13), an effect attributed to the activity of the β-keto acyl-ACP synthase FabF (14, 15). How transcriptional regulation and the temperature sensitivity of fatty acid synthesis each contribute to homeoviscous adaptation is unclear.

Here, we use a systems approach to determine how *E. coli* achieves homeoviscous adaptation. By comprehensively quantifying enzymes and metabolites of the fatty acid and phospholipid synthesis pathways, we reveal how *E. coli* integrates a temperature-sensitive metabolic valve with a transcriptional feedback loop to accelerate response times and adapt membrane composition within a single cell cycle. We use a quantitative model to determine how the system is calibrated to produce specific membrane compositions for each temperature. We predict that homeoviscous adaptation systems in diverse organisms will feature similar regulatory motifs that accelerate adaptation to temperature shocks.

## Results

### Steady-state fatty acid synthesis enzyme concentrations exhibit paradoxical correlations with membrane lipid saturation

*E. coli* fatty acids are produced as thioesters attached to an acyl carrier protein (acyl-ACP) by the saturated and unsaturated branches of the fatty acid synthesis pathway **(Figure 1B, Supplemental Figure 1)**. The first enzyme of the *E. coli* phospholipid synthesis pathway (PlsB) transfers a C16:0 fatty acid (saturated) or a C18:1 fatty acid (unsaturated) from the corresponding acyl-ACP to the *sn-* 1 position of glycerol-3-phosphate (G3P), producing lysophosphatidic acid (LPA). Next, the enzyme PlsC transfers a fatty acid from either C16:1 or C18:1 ACP to the *sn-*2 position of LPA to generate phosphatidic acid (PA). Because PlsC chiefly transfers unsaturated fatty acyl chains whereas PlsB transfers both saturated and unsaturated fatty acyl chains, membrane fluidity is primarily determined by the fatty acid incorporated at *sn-*1 by PlsB.

We prepared cultures of *E. coli* NCM3722 using defined minimal medium (MOPS/0.2% glycerol) at 5 temperatures from 12 to 42 °C. Exponential-phase cultures (OD600 = 0.3-0.4) were rapidly sampled and quenched to preserve steady-state metabolite concentrations (16). Proteins and phospholipids were extracted and quantified using liquid chromatography/mass spectrometry (LCMS) (17). As expected, the abundance of unsaturated fatty acyl chains within phospholipid intermediates and membrane phospholipids decreased with increasing temperature **(Figure 1D)**. Remarkably, the proportions of phospholipids with saturated or unsaturated *sn*-1 acyl chains closely correspond to the proportions of saturated and unsaturated acyl-ACP substrates of PlsB: saturated acyl-ACP (C16:0 ACP) and 16:0 *sn*-1 phospholipids increase with temperature, while unsaturated acyl-ACP (C18:1 ACP) and 18:1 *sn*-1 phospholipids decrease. Likewise, the proportions of 16:1 and 18:1 *sn*-2 phospholipids also closely match the PlsC acyl-ACP substrate pool, with the abundance of 16:0 *sn-*2 phospholipids increasing with temperature despite inefficient utilization of C16:0 ACP by PlsC (18) **(Figure 1D, Supplemental Figure 2A & 2B)**. These measurements confirm that the composition of phospholipid acyl chains is determined by the compositions of the PlsB and PlsC substrate pools.

Steady-state concentrations of C16:0 and C18:1 ACP may be determined by the enzymes that split the fatty acid synthesis pathway into saturated and unsaturated branches (FabI, FabA, and FabB). Expression of FabA and FabB is controlled by the transcriptional regulators FabR and FadR. FabR regulates expression of its targets according to the ratio of C16:0 ACP to C18:1 ACP (12), whereas FadR primarily regulates fatty acid catabolism in response to acyl-CoA thioesters synthesized from exogenous fatty acids (19) **(Figure 1C)**. To determine how transcriptional regulation controls the composition of the acyl-ACP substrate pool, we quantified fatty acid pathway enzymes at each temperature. While concentrations of most enzymes (including FabI) remained relatively constant **(Supplemental Figure 2C)**, FabA concentrations decreased ∼2-fold from 12 to 42 °C, while FabB concentrations increased ∼2-fold **(Figure 1E)**. This trend contradicts expectations as multiple studies report that increasing FabB or decreasing the FabA/FabB ratio at constant temperature increases unsaturated fatty acid production at the expense of saturated fatty acids (20–22). We confirmed that FabA overexpression increases the proportion of saturated phospholipids, while FabB overexpression increases the proportion of unsaturated phospholipids **(Figure 1F)**.

### The fatty acid pathway exhibits overshoot kinetics following temperature shocks

The unexpected relationship between membrane composition and the steady-state FabA/FabB ratio suggests that factors in addition to transcriptional regulation determine the composition of the PlsB and PlsC substrate pools. The temperature sensitivities of enzymes within the fatty acid pathway increase the production of unsaturated fatty acids at cold temperatures, an intriguing form of post-translational regulation of metabolism by temperature thought to be enabled by FabF (13). To better resolve how the temperature sensitivities of fatty acid synthesis enzymes contribute to thermal adaptation, we tracked acyl-ACP and phospholipids in cultures subject to a rapid cold shock from 37 to 13 °C **(Figure 2A)**. The cold shock immediately perturbed both acyl-ACP and phospholipid intermediate pools and arrested growth for approximately one hour **(Supplemental Figure 3, Figures 2B, 2C)**. Strikingly, within 5 minutes of the cold shock C16:0 ACP decreased by approximately 5-fold, while C18:1 ACP remained comparatively stable **(Supplemental Figure 3)**.

**Figure 2.**
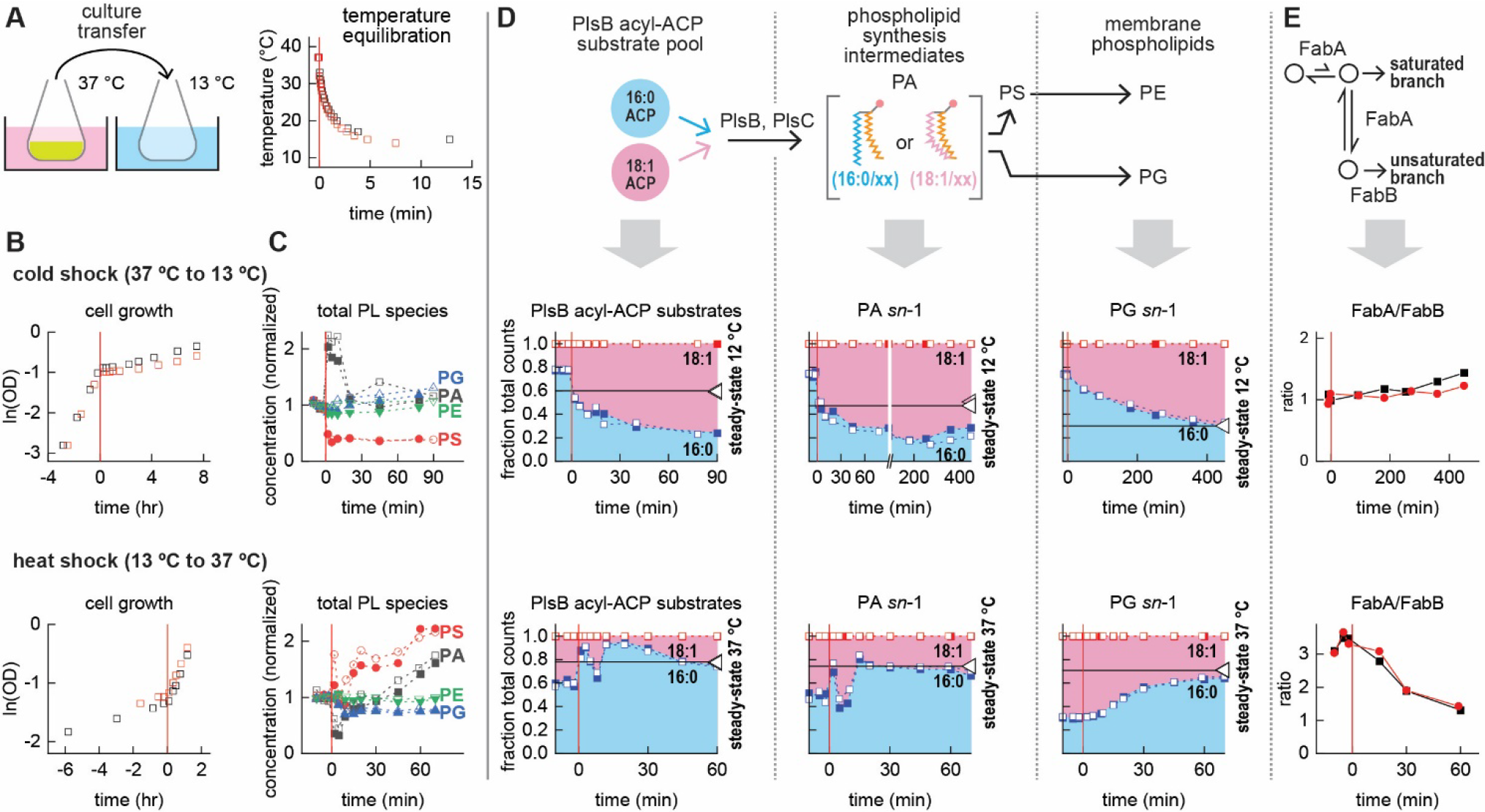
The fatty acid synthesis pathway responds immediately to temperature shocks with overshoot kinetics. **A.** Exponential-phase cultures are transferred to empty flasks pre-equilibrated in a temperature-controlled bath. **B, C.** Effects of temperature shocks on growth **(B)** and phospholipid abundance **(C)**. Total quantities of PA, phosphatidylserine (PS), PE, and PG normalized to average of their pre-shift values. **D.** Compositions of PlsB substrate, PA, and PG pools before and after temperature shocks (indicated with vertical red lines at 0 min). Final steady-state values indicated by horizontal lines to highlight the overshoot. **E.** FabA/FabB ratios following cold and heat shocks. All graphs depict kinetic series obtained from 2 independent experiments.

These changes inverted the relative proportions of saturated and unsaturated fatty acid substrates of the PlsB substrate pool, making C18:1 ACP the most abundant PlsB substrate **(Figure 2D)**. In parallel with the acyl-ACP pool, the phospholipid intermediate pool also rapidly changed: 18:1 *sn*-1 PA considerably increased at the expense of 16:0 *sn-*1 PA **(Figure 2D, Supplemental Figure 4)**. As these dynamics are too rapid to be mediated by transcription, the changes observed must result from the direct influence of temperature upon the fatty acid synthesis pathway (13).

A close examination reveals that the size of the C18:1 ACP fraction immediately following the cold shock far exceeds the fraction observed during steady-state growth at 12 °C (indicated by the horizontal lines in **Figure 2D)**. This overshoot is also reflected in the excess proportion of 18:1 *sn*-1 PA, which persists for at least 7 hours following the cold shock. Remarkably, the 18:1 *sn*-1 PG fraction reaches its final steady-state 12 °C level within 8 hours, approximately one doubling period during steady-state growth (7 hours) **(Figure 2D)**.

We quantified fatty acid synthesis enzymes to identify how transcriptional regulation contributes to the cold shock response. Within the 7.5 hour period monitored following the cold shock, all enzyme concentrations remain stable aside from FabB, which begins to decrease after 5 hours, thus increasing the FabA/FabB ratio **(Figure 2E, Supplemental Figure 5)**. As the FabA/FabB ratio does not substantially change for up to 7.5 hours after the cold shock, the adaptations observed are achieved entirely by the direct influence of temperature on fatty acid synthesis enzymes.

To determine how the fatty acid pathway responds to increasing temperature, we analysed fatty acid and phospholipid synthesis intermediates after a heat shock (rapid transfer from 13 °C to 37 °C). Cell growth immediately accelerated after the heat shock **(Figure 2B)**. The fatty acid and phospholipid intermediate pools responded within 1 minute and exhibited behaviours consistent with accelerated phospholipid synthesis **(Figure 2C)**. Remarkably, most fatty acid intermediates following the heat shock follow trajectories that completely reverse those observed following the cold shock **(Supplemental Figure 3)**. Notably, C16:0 ACP increases more than 2-fold while C18:1 ACP decreases sharply; as a result, C16:0 ACP becomes the most abundant PlsB substrate within 2 minutes of the heat shock, which in turn increases the C16:0 *sn-*1 PA and PG fractions **(Figure 2D)**. Similarly to the behaviour of the C18:1 ACP fraction following cold shocks, the C16:0 ACP fraction of the PlsB substrate pool briefly overshoots the steady-state value at 37 °C. The C16:0 *sn-* 1 fraction of the membrane phospholipid PG closely approaches the final steady-state levels within one doubling period (58 minutes).

We quantified enzymes of the fatty acid synthesis pathway to determine how transcriptional regulation contributes to the heat shock response. Aside from FabA and FabB, the concentrations of all enzymes monitored remained stable **(Supplemental Figure 5)**. FabA and FabB did not substantially change within 15 minutes of the heat shock, indicating that the initial dynamics are not driven by transcriptional regulation. FabB concentrations increased after 15 minutes and reached ∼2-fold of the pre-shock level after 60 minutes, whereas FabA decreased slightly. As a consequence, the FabA/FabB ratio decreased ∼2-fold, nearly reaching the ratio observed in steady-state conditions at 37 °C **(Figure 2E)**. Given that FabB overexpression increases unsaturated fatty acid synthesis, the transcriptional response that increases FabB expression increases unsaturated fatty acid synthesis and corrects the excess synthesis of saturated fatty acids that is caused by the temperature sensitivities of fatty acid synthesis enzymes. Therefore, the apparent paradox of FabA and FabB concentrations observed during steady-state growth can be understood as partially counteracting the direct influence of temperature upon the fatty acid pathway.

### Temperature shifts reroute fatty acid flux between saturated and unsaturated branches by tuning relative activities of FabB and FabI

Our comprehensive measurements of fatty acid intermediates allow us to evaluate mechanisms by which temperature directly controls the output of the *E. coli* fatty acid pathway. Temperature-induced changes in membrane lipid composition are proposed to be mediated by the β-keto-acyl-ACP synthase FabF, which extends acyl chains throughout the fatty acid pathway with the exception of C10:1-3*Z* ACP **(Supplemental Figure 1)**. FabF is essential for membrane viscosity control as it initiates synthesis of C18:1 ACP by elongating C16:1 ACP (14, 23). *E. coli* Δ*fabF* cannot synthesize sufficient C18:1 ACP and therefore is unable to adapt membrane lipids to temperature (24). The activity of FabF remains relatively stable with decreasing temperature, a property suggested to allow cold temperatures to directly increase synthesis of C18:1 ACP and thus the proportion of 18:1 *sn-*1 phospholipids in *E. coli* (14, 15). A superficial examination of our temperature shift data supports this hypothesis: cold shocks deplete C16:1 ACP and increase the intermediate C18:1-OH ACP, suggesting increased elongation by FabF **(Figure 3A)**. Conversely, heat shocks decrease both C18:1-OH ACP and C18:1 ACP, indicating reduced C16:1 ACP elongation by FabF. However, cold shocks and heat shocks also trigger profound changes in C16:0 ACP concentration **(Figure 3B)**, which also determines PlsB substrate pool composition and membrane fluidity. How concentrations of C16:0 ACP and other fatty acid synthesis intermediates might be affected by FabF activity is not clear.

**Figure 3.**
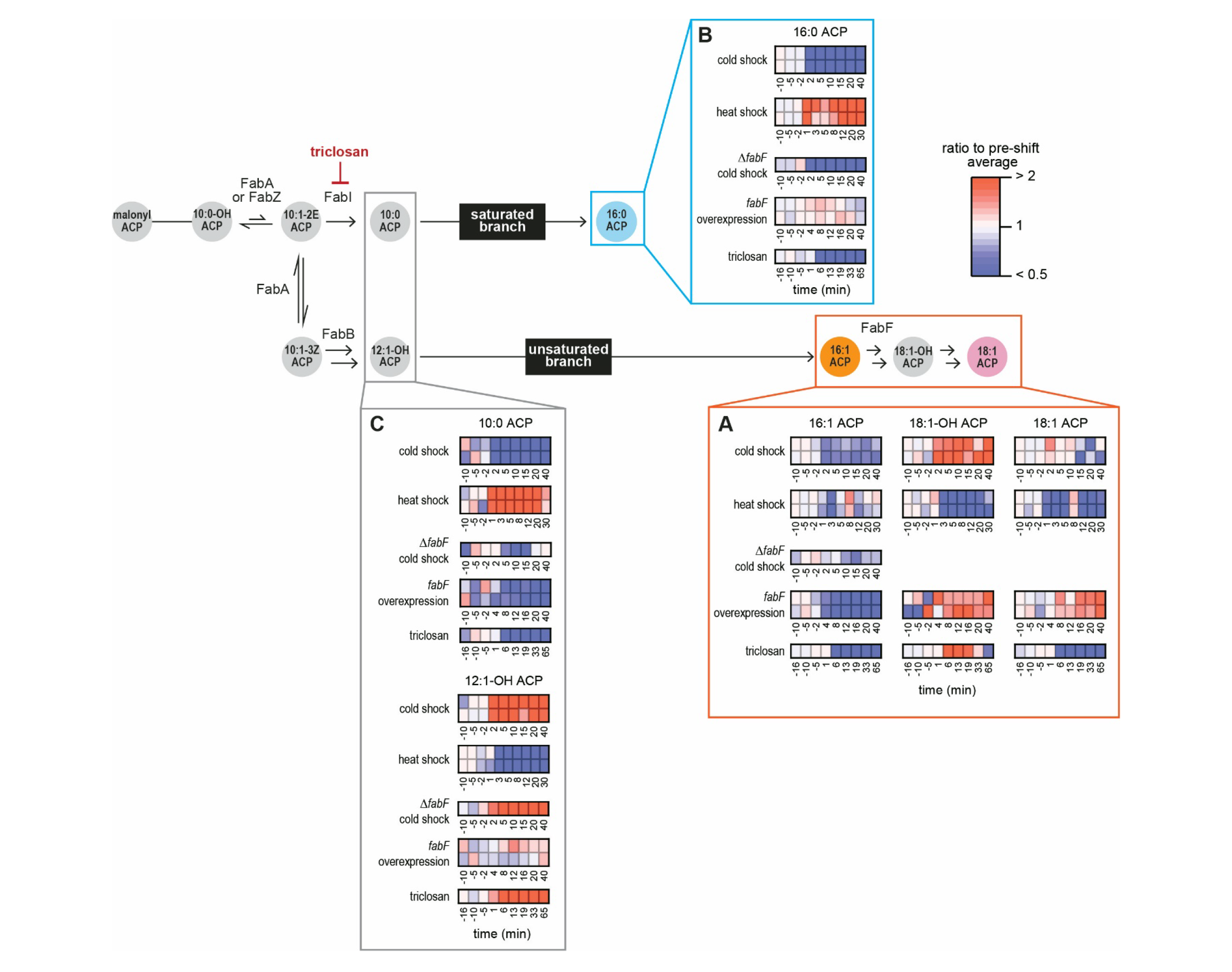
Comparison of acyl-ACP dynamics following temperature shocks, cold shock in a Δ*fabF* strain, *fabF* overexpression in wild-type *E. coli*, and FabI inhibition with triclosan. Values for each time point are normalized by the average value obtained from the 3 measurements preceding the perturbation. **A & B.** Products of the unsaturated **(A)** and saturated branches **(B). C.** Concentrations of FabI and FabB products immediately following the saturated-unsaturated pathway branch point.

We tested whether FabF initiates temperature responses by subjecting a *fabF* deletion mutant (*E. coli* Δ*fabF*) to a cold shock. As expected, *E. coli* Δ*fabF* produced only trace amounts of C18:1 ACP before and after the cold shock **(Supplementary Figure 6A)** (15). However, C16:0 ACP decreased by 2-fold following the cold shock in *E. coli* Δ*fabF*, a response also observed in wild-type *E. coli* **(Figure 3B)**. Furthermore, the cold shock dynamics of intermediates throughout the fatty acid synthesis pathway are highly similar in both wild-type *E. coli* and Δ*fabF* strains: specifically, acyl-ACP products of FabI decreased, while hydroxyl-acyl-ACP species in the unsaturated branch increased by more than 2-fold **(Supplementary Figure 6B)**. To test whether increased FabF activity depletes C16:0 ACP, we placed *fabF* under control of an inducible promoter and monitored acyl-ACP dynamics immediately following *fabF* induction. While *fabF* overexpression increased C18:1- OH ACP and C18:1 ACP at the expense of C16:1 ACP **(Figure 3A)**, C16:0 ACP concentrations did not decrease **(Figure 3B)**. These results strongly suggest that while FabF is required to produce C18:1 ACP and thereby enable PlsB to incorporate unsaturated fatty acids into phospholipids, FabF itself does not mediate temperature-dependent flux allocation between saturated and unsaturated pathways.

Flux partitioning between two branches of a metabolic pathway is determined by the relative activities of the enzymes catalysing the committed steps. We therefore considered instead whether the enzymes that catalyse the reactions that commit fatty acid intermediates to the saturated and unsaturated branches (FabI and FabB) might respond to temperature and thereby adapt the proportions of saturated and unsaturated fatty acids. As cold shocks tend to decrease the flux channelled into the saturated branch via FabI, we reasoned that a similar effect may be obtained by directly inhibiting FabI. We therefore monitored the response of the fatty acid synthesis pathway immediately after adding the FabI inhibitor triclosan (25). Triclosan caused most hydroxyl- and 2- enoyl-acyl-ACP species to accumulate at the expense of acyl-ACP **(Supplemental Figure 7)**.

Similarly to the cold shock, triclosan decreased C16:0 ACP and C16:1 ACP and increased C18:1- OH ACP **(Figure 3A & 3B)**. Interestingly, the dynamics of most fatty acid intermediates following FabI inhibition closely resembled the cold shock response **(Supplemental Figure 6B)**. Both triclosan and cold shock decreased the acyl-ACP products of FabI throughout the pathway.

Interestingly, both treatments decreased concentrations of hydroxyl-acyl-ACP intermediates within the saturated branch but increased hydroxyl-acyl-ACP concentrations within the unsaturated branch **(Supplemental Figures 6B&7)**. Furthermore, both treatments decrease the FabI product C10:0 ACP while increasing the first detected intermediate of the unsaturated pathway (C12:1-OH ACP). These dynamics are observed in the cold shock responses of both wild-type and *E. coli* Δ*fabF* **(Figure 3C)**. In contrast to the cold shock, *fabF* overexpression does not increase C12:1-OH ACP. Importantly, the cold shock responses of branch point products are exactly reversed following heat shock: concentrations of FabI products increase throughout the pathway, whereas hydroxyl-acyl-ACP concentrations decrease, consistent with an increase in FabI activity relative to other pathway enzymes. These similarities suggest that temperature allocates flux between saturated and unsaturated fatty acid synthesis by tuning the balance between FabI and FabB activities, thereby varying the proportions of C16:0 ACP and C18:1 ACP.

### A simple mathematical model recapitulates temperature dependence of steady-state membrane composition

Our steady-state and temperature shock experiments reveal two components that contribute to homeoviscous adaptation: 1) the direct influence of temperature on the enzymes at the branch point between saturated and unsaturated fatty acid pathways, and 2) a transcriptional response that shifts FabB expression to partially counteract the direct influence of temperature upon the fatty acyl-ACP pool. The delayed increase in FabB expression following heat shock **(Figure 2E)** and the low steady-state FabB expression at cold temperatures **(Figure 1E)** are both consistent with increasing FabR repression of FabB expression due to C18:1 ACP abundance (12). To test whether temperature sensitivity and regulation by FabR are sufficient to reproduce our observations and thereby account for the *E. coli* homeoviscous adaptation response, we built a differential equation-based model that emulates a temperature-controlled branched pathway with a transcriptional feedback loop. The simulated branched pathway consists of two enzymes (FabI and FabB) that produce saturated and unsaturated precursors, respectively (representing 16:0 and 18:1 acyl-ACP). These precursors are converted to corresponding membrane phospholipids by an enzyme operating at constant flux (analogous to PlsB) **(Figure 4A)**. To simulate transcriptional regulation, we set FabI expression at a constant rate while placing FabB expression under control of a FabR- mediated negative feedback loop that represses FabB in proportion to the abundance of the unsaturated precursor **(Figure 4B).** To maintain steady-state concentrations, all species included within the model (enzymes, precursors, and membrane phospholipids) are steadily depleted at a constant rate *μ* to simulate continuous dilution by cell growth.

**Figure 4.**
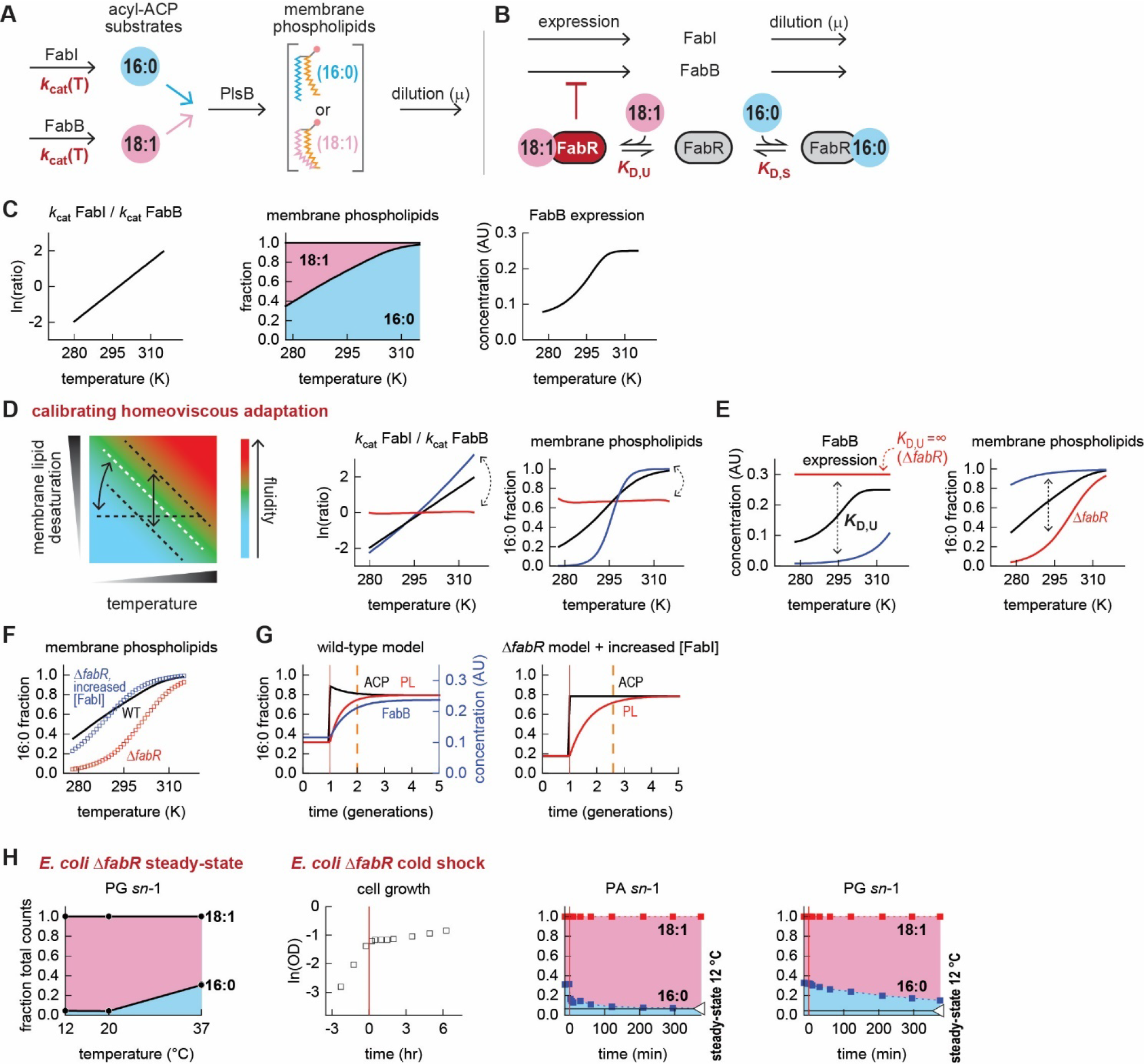
A mathematical model recapitulates all core behaviours of the *E. coli* homeoviscous adaptation system. **A.** Simulated phospholipid synthesis from an acyl-ACP pool consisting of saturated (16:0) and unsaturated acyl-ACP (18:1). **B.** Simulated regulation of FabB expression by FabR. **C.** Asymmetric temperature sensitivities of FabB and FabI increases both the saturated proportion of membrane phospholipids and the abundance of FabB in parallel with temperature. **D.** Exploring how model parameters calibrate the homeoviscous system to produce desired membrane lipid desaturation at different temperatures. Varying the temperature sensitivities of FabB and FabI varies how the fractions of membrane phospholipids change with temperature. **E.** Varying the FabR- 18:1 ACP complex dissociation constant changes the relationship between temperature and FabB expression, which in turn affects the steady-state saturated phospholipid fraction while retaining the overall temperature dependency of membrane lipid saturation. **F.** Varying the basal expression of FabI enables a Δ*fabR* model to match the membrane composition of a model that retains FabR control (wild-type). **G.** Dynamic response to heat shock generated by wild-type model exhibits overshoot kinetics and accelerates adaptation of the membrane phospholipid pool, achieving 90% adaptation within 1 generation (indicated by dashed orange line) The Δ*fabR* model does not exhibit overshoot kinetics and requires 1.6 generations to reach steady-state membrane composition. **H.** Cold shock of a Δ*fabR* knockout confirms model predictions: without transcriptional repression by FabR, no overshoot kinetics are possible and homeoviscous adaptation requires more than one generation time. Steady-state data from three samples obtained from one culture prepared at each temperature. Kinetic series obtained from one experiment.

Guided by our evidence indicating that cold temperatures reroute flux by decreasing FabI activity relative to FabB, we modelled asymmetric temperature dependencies in the FabI and FabB *k*cat values such that FabI *k*cat decreases more rapidly at low temperatures than FabB. This asymmetry causes the steady-state concentration of saturated membrane phospholipids to increase with temperature **(Figure 4C)**. As also observed in our experiments, increasing temperatures decrease the abundance of the unsaturated precursor, which relieves FabB repression by FabR and allows FabB abundance to increase. Thus, our model captures the essential steady-state behaviours of homeoviscous adaptation. Next, we tuned the parameters of our model to explore how the *E. coli* homeoviscous adaptation system might calibrate the composition of the fatty acid precursor pool across temperatures. Varying the temperature sensitivity of FabI *k*cat changed the temperature sensitivity of the FabI/FabB *k*cat ratio, which in turn adjusted the sensitivity of phospholipid composition to temperature **(Figure 4D)**. The membrane composition remains unchanged across temperatures if FabI and FabB *k*cat temperature sensitivities are identical. Therefore, an asymmetric temperature dependence in the activities of FabI and FabB is both necessary and sufficient to cause membrane composition to vary with temperature.

We next sought to determine how transcriptional regulation by FabR contributes to steady-state membrane composition. Tuning the affinity of the FabR regulator for the unsaturated acyl-ACP precursor shifted the temperature sensitivity of FabB abundance and the saturated phospholipid fraction **(Figure 4E)**. Eliminating the interaction between FabR and the unsaturated precursor (equivalent to a Δ*fabR* strain) abolishes the temperature sensitivity of FabB expression, which remains at maximum concentration. This “Δ*fabR* model” nevertheless retains temperature control of membrane composition but increases the fraction of unsaturated phospholipids at all temperatures **(Figure 4E)**. Therefore, both transcriptional regulation parameters and the temperature sensitivities of FabI and FabB may be calibrated by evolution to produce specific membrane compositions across a range of temperatures.

### Transcriptional regulation by FabR is required to accelerate homeoviscous adaptation

Models lacking FabR regulation retain temperature control of membrane composition but generate far more unsaturated membrane phospholipids at all temperatures than models with otherwise identical parameters. However, such Δ*fabR* models nevertheless recover the membrane compositions of the parent “wild-type” models at all temperatures if the basal expression level of FabI is increased **(Figure 4F)**. This raises questions about the utility of transcriptional regulation by FabR: why does *E. coli* retain transcriptional regulation by FabR when a temperature-sensitive branch point is sufficient to adapt membrane composition?

The delayed effects of temperature shocks on FabB expression observed in our experiments suggest a role for transcriptional regulation. The immediate homeoviscous response driven by temperature control of the FabB/FabI activity ratio initiates membrane adaptations by producing unsaturated (or saturated) fatty acids at proportions that far exceed final steady-state proportions.

Over time, transcriptional regulation partially counteracts the overshoot by shifting FabB expression to steer the acyl-ACP pool composition to its final steady-state. However the initial excess production of saturated or unsaturated fatty acids accelerates thermal adaptation and enables *E. coli* to quickly adapt its membrane to new temperatures. As transcriptional regulation is used to correct the initial overshoot following temperature shocks, FabR may be retained to enable the overshoot kinetics required to accelerate membrane adaptation.

We first tested whether a “wild-type” model (featuring both temperature regulation of FabI/FabB activity and FabB repression by FabR) could exhibit overshoot kinetics and achieve the rapid membrane adaptation observed in our experiments. We modelled temperature shocks and tracked the composition of the acyl-ACP and membrane phospholipid pools. In response to a heat shock, the saturated acyl-ACP fraction initially surpassed the final steady-state value, mimicking the experimentally-observed overshoot **(Figure 4G)**. The overshoot in saturated acyl-ACP abundance decreased repression of FabB expression by FabR, causing steady-state FabB concentration to increase. Increasing FabB in turn increased unsaturated precursor synthesis, which corrected the overshoot and steered the composition of the acyl-ACP pool to its final steady-state. Furthermore, the membrane phospholipid pool reached within 90% of its final composition within 1 generation following the heat shock.

We next simulated the thermal response of a pathway retaining differential temperature regulation of FabI/FabB activity but lacking transcriptional regulation by FabR (Δ*fabR* model). Subjecting the Δ*fabR* model to a simulated heat shock immediately shifted the precursor pool to its final steady-state composition **(Figure 4G).** No overshoot of the acyl-ACP pool was observed as FabB abundance could not respond to changes in the 18:1 precursor. Furthermore, despite retaining the immediate response of the temperature-sensitive branch point, the Δ*fabR* model required 1.6 generation times to reach 90% of its final steady-state composition at the new temperature.

We tested our model by constructing an *E. coli* Δ*fabR* strain and characterized the steady-state membrane composition of cultures prepared at 3 temperatures. As previously observed (26) and as replicated in our model, the unsaturated proportion of membrane phospholipids increased in *E. coli* Δ*fabR* at all temperatures compared to wild-type **(Figure 4H)**. However, *E. coli* Δ*fabR* exhibited higher proportions of unsaturated phospholipids at low temperatures, confirming that FabR is not required for membrane adaptation. To test whether FabR is necessary for overshoot kinetics as predicted by our model, we subjected *E. coli* Δ*fabR* to a cold shock. The cold shock immediately increased the proportion of 18:1 *sn*-1 PA intermediates to its final steady-state values without an overshoot **(Figure 4H)**. Consistent with our model results, the proportion of 18:1 *sn*-1 PG did not reach its final steady-state value within one generation time (7.5 hours). This confirms that FabR enables the fatty acid pathway to overshoot the final steady-state composition following temperature shocks, which in turn reduces the time required to adapt the membrane composition to new temperatures. In other words, the role of FabR is not to enable homeoviscous adaptation but to accelerate homeoviscous adaptation.

### Rapid homeoviscous adaptation accelerates growth recovery following cold shocks in respiration-dependent medium

The evolution of a transcriptional regulator that accelerates homeoviscous adaptation implies that the ability to rapidly restore optimal fluidity is advantageous. However, the benefits of rapid homeoviscous adaptation have proven elusive in experimental settings. While extreme perturbations to membrane composition have long been known to compromise viability, artificial perturbations that remain within the typical range of thermal adaptation do not exert an obvious effect (27–29), suggesting that bacteria readily tolerate moderate alterations in membrane fluidity. For example, *E. coli* Δ*fabF* is unable to vary membrane composition in response to temperature, but grows robustly at low temperatures (30) and recovers growth following cold shocks as quickly as wild-type strains (24). Consistent with these observations, our *E. coli* Δ*fabR* strain grew normally following a cold shock **(Figure 4H)**. These findings raise the question of how the rapid homeoviscous adaptation response provides an apparent evolutionary advantage to *E. coli* and other organisms.

A recent discovery revealed that respiratory metabolism is highly sensitive to membrane fluidity (21). This is due to the dependence of respiration upon electron transport reactions between membrane-bound oxidoreductases and electron carrier substrates. As diffusion of membrane-bound reactants is highly dependent upon membrane viscosity, even incremental perturbations in membrane composition impact growth when catabolism depends upon respiration. For example, the first step of succinate catabolism (oxidation of succinate to fumarate) requires reduction of a membrane-bound ubiquinone that must be subsequently recycled by an oxidase. To test whether homeoviscous adaptation is required for optimal steady-state growth in respiration-dependent conditions, we compared growth of wild-type *E. coli* against *E. coli* Δ*fabF* in three defined media: rich glucose medium with casamino acids, glycerol minimal medium, and succinate minimal medium. Consistent with prior observations, *E. coli* Δ*fabF* grew normally in both glucose and glycerol medium, but exhibited severely reduced growth in succinate medium **(Figure 5A)**. Growth of *E. coil* Δ*fabR* is not impaired in succinate medium likely because its membrane fluidity is not decreased relative to wild-type, as proportions of unsaturated membrane lipids in *E. coil* Δ*fabR* exceed wild-type levels **(Figure 4H)**. To confirm that the growth defect of the Δ*fabF* strain is caused by low levels of unsaturated phospholipids, we added the fatty acid *cis-*vaccenate, which is converted to C18:1-CoA and used as a substrate by both PlsB and PlsC. *Cis-*vaccenate completely restored growth of the Δ*fabF* strain to match the wild-type strain **(Figure 5A)**. As *cis-*vaccenate did not increase growth of the wild-type strain, the growth recovery of *E. coli* Δ*fabF* is likely mediated by restored membrane fluidity due to incorporation of C18:1 acyl chains into membrane phospholipids.

**Figure 5.**
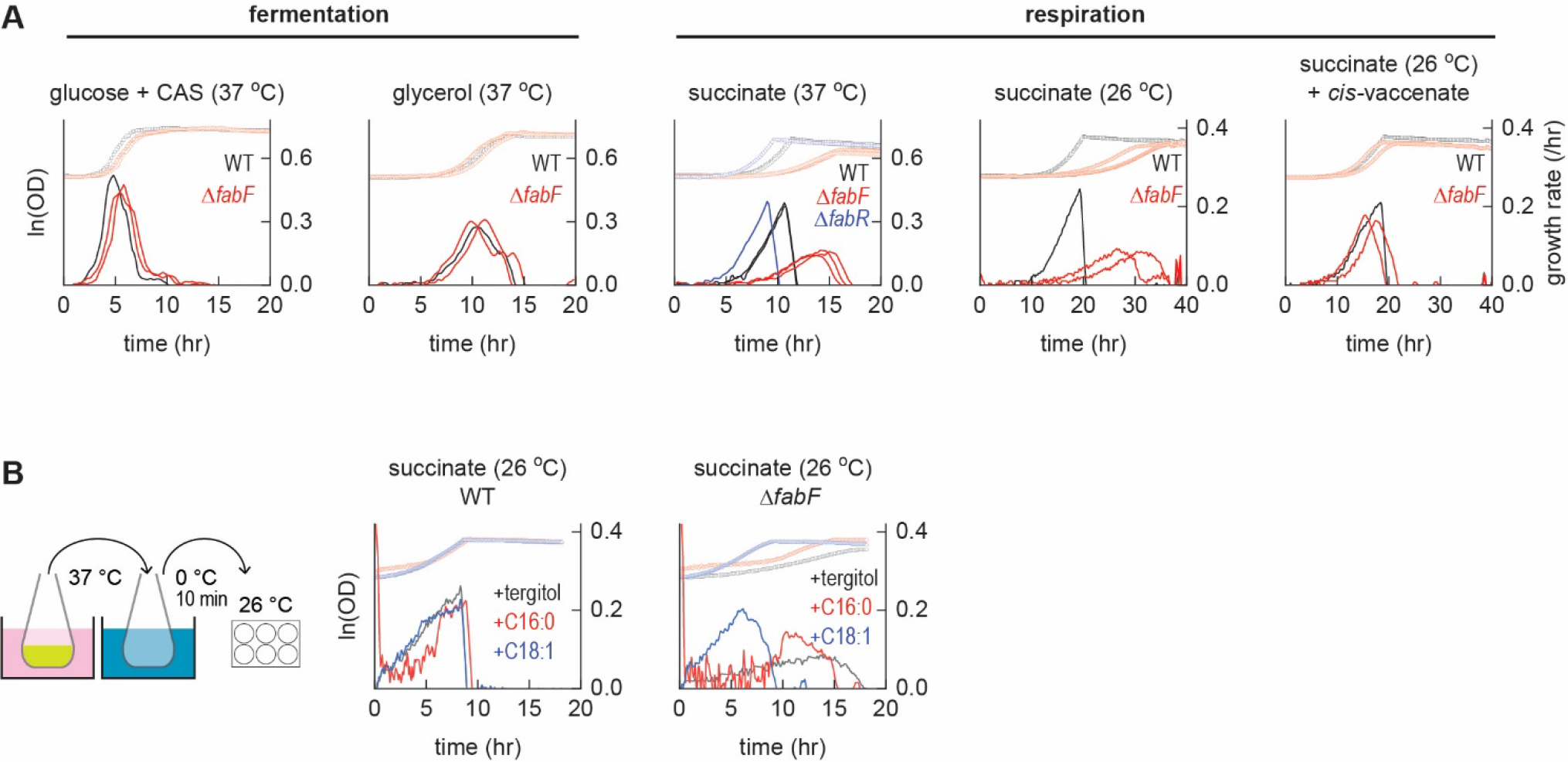
Increased synthesis of unsaturated phospholipids accelerates growth recovery from a cold shock in respiration-dependent conditions. Growth of cultures monitored in a 6-well plate by optical density measurements (symbols). Derivatives of the growth curves (smoothed over 7 points) yields instantaneous growth rates (lines); each line represents an independent culture. **A.** Comparison of growth of wild-type (WT), Δ*fabR,* and Δ*fabF* strains in MOPS-based defined media containing indicated carbon sources at indicated temperatures (0.2% w/vol glucose, glycerol, or succinate; glucose media also contains 0.1% w/vol CAS amino acids). **B.** Comparison of wild-type (WT) and Δ*fabF* strains following a cold shock from 37 °C to 0 °C followed by recovery in a microplate reader at 26 °C with and without supplementation with tergitol carrier, palmitate (C16:0), or *cis-*vaccenate (18:1).

We next tested whether the rapid adaptation of membrane fluidity accelerates recovery of growth following a cold shock. Therefore, we tested whether growth recovery in *E. coli* Δ*fabF* could be accelerated by addition of *cis-*vaccenate. Wild-type and Δ*fabF* succinate cultures prepared at 37 °C were transferred during exponential phase to an ice bath and incubated for 10 minutes before culturing at 26 °C. As expected, Δ*fabF* cultures grew extremely slowly following the cold shock unless supplemented with *cis-*vaccenate, which enabled immediate growth recovery and steady-state growth rate that matched the wild-type rate **(Figure 5B)**. The addition of the saturated fatty acid palmitate did not recover growth of the Δ*fabF* culture, indicating that the growth recovery enabled by *cis-*vaccenate is due to restoration of unsaturated phospholipid biosynthesis and membrane fluidity rather than consumption of the fatty acid.

## Discussion

The broad conservation of homeoviscous adaptation indicates that the ability to precisely maintain membrane fluidity by incrementally adjusting membrane composition is an essential trait in natural environments. How organisms calibrate membrane composition in response to temperature is poorly understood. Several aspects of the homeoviscous adaptation system in *E. coli* have been identified, including the control of membrane composition by the acyl-ACP pool (11) and direct temperature control of unsaturated fatty acid synthesis via a transcription-independent mechanism (13), while the role of transcriptional regulation by FabR has remained obscure (12, 26). Our study advances the field by revealing how these components are integrated into a system that measures temperature, adapts fatty acid synthesis, and rapidly restores membrane fluidity. These insights are made possible by combining comprehensive measurements of metabolites and enzymes of the fatty acid and phospholipid pathways with a quantitative model that reproduces the core properties of homeoviscous adaptation in *E. coli*. We find that *E. coli* adjusts membrane composition by using a fatty acid pathway that partitions flux between its saturated and unsaturated branches according to temperature. In this system, the temperature sensitivity of the fatty acid branch point is the most basic element required for homeoviscous adaptation, which links the supply of unsaturated fatty acids to requirements for membrane fluidity in cold temperatures. Our data indicate that the temperature sensitivity of unsaturated fatty acid synthesis most likely arises from properties of the enzymes that commit fatty acid intermediates to the saturated and unsaturated fatty acid pathways (FabB and FabI), rather than from properties of FabF. Our model suggests the temperature-dependent activities and expression levels of FabB and FabI may be calibrated by evolution to produce specific membrane compositions that stabilize membrane viscosity at any growth temperature.

The implementation of temperature control within the fatty acid pathway rather than a hypothetical temperature-sensitive transcriptional regulator ensures an immediate response to temperature shocks. This acceleration is particularly important because *E. coli* is unable to adjust fluidity by modifying its membrane lipids. Therefore, restoring membrane fluidity requires substantially diluting the ∼10^7^ phospholipids that comprise its membrane with phospholipids newly-assembled from an adapted fatty acid pool. However, a temperature-sensitive metabolic pathway is insufficient on its own to adapt the membrane to new temperatures within a single generation.

Counterintuitively, the accelerated adaptation we observe is made possible by a slow-responding feedback loop implemented by FabR, a transcriptional regulator whose role has remained obscure (12, 26, 31). Negative autoregulation of the unsaturated fatty acid pathway allows the initial temperature shock response to overshoot the eventual steady-state target. Changes in C18:1 ACP concentration lead FabR to correct the overshoot by adjusting FabB expression. Importantly, steady-state FabB concentrations are set at all temperatures to ensure that cold and heat shocks will always generate an overshoot and thereby accelerate homeoviscous adaptation. The combination of two regulatory components that respond at different timescales is a recurring biological motif that accelerates adaptation (32).

We hypothesize that the general principles revealed here exist in any microorganism that is capable of rapid homeoviscous adaptation. Specifically, we predict that all homeoviscous adaptation systems will feature networks that enable overshoot kinetics. Overshoot kinetics can be achieved by protein degradation, negative autoregulation, or incoherent feedforward loops. Responses consistent with overshoot kinetics have been observed in prokaryotes with Des-type membrane fluidity sensors (*Bacillus megaterium and B. subtilis*) (33–35), and in multicellular eukaryotes (6), which use systems very different from *E. coli*. Membrane fluidity in the food pathogen *Listeria monocytogenes* is modulated by tuning the branching position of fatty acids (36). The abundance of fluidity-enhancing branched-chain fatty acids is at least partly controlled by the selection of the CoA primer by the *L. monocytogenes* FabH homologue, preference for which is sensitive to temperature (37). This provides a possible mechanism for initiating rapid adaptation of the *L. monocytogenes* membrane analogous to the temperature sensitivity of the FabI/FabB branch point in the *E. coli* pathway. Mathematical modelling of the *L. monocytogenes* fatty acid pathway based upon FabH substrate selectivity alone was unable to reproduce measurements of anteiso-branched phospholipids, leading the authors of the work to suggest an additional step required to attain the final composition attained (38). Our findings suggest that *L. monocytogenes* homeoviscous adaptation may involve a slow-responding regulatory component that accelerates adaptation time (analogous to FabR) yet to be discovered.

## Acknowledgements

We thank Leander Lutze, Professor Sarah L. Keller, and the Systems Biology Department at VU Amsterdam for insightful discussions. Project supported by start-up funds to GB from the TU Delft Bionanoscience Department.

## Author contributions

LH: designed and performed steady-state temperature experiments and pilot temperature experiments, evaluated data. SvdB: built data analysis pipeline, designed and performed triclosan experiments, evaluated data. EG: cloned fatty acid synthesis enzymes, performed initial *fabF* overexpression experiments. FY, AZ-D: LCMS analysis. FB: mathematical modelling. GB: supervised project, performed temperature shift, microplate cultures, and overexpression experiments, evaluated data, and wrote the manuscript. All authors approved the final manuscript.

## Competing interests

The authors declare no competing interests.

## Materials and Methods

### Culture conditions

Unless otherwise indicated, cultures were grown in Erlenmeyer flasks with 0.2% glycerol / MOPS minimal medium with 0.2% (w/v) glycerol (39). Flasks were incubated in a water bath (Grant Instruments Sub Aqua Pro) and stirred with a magnetic bar (1200 rpm) coupled to a magnetic plate (2mag MIXdrive 1 Eco and MIXcontrol 20). Optical density was measured using Ultrospec 10 Cell Density Meter (GE Healthcare). Fatty acid solutions (40 mM) were prepared by solubilisation in 26% tergitol and neutralization to pH 7 using NaOH. For temperature shifts, 25-35 mL medium was transferred to an empty Erlenmeyer flask incubated in a water bath.

### Plasmids and strains

*Escherichia coli* K-12 strain NCM3722 (CGSC #12355) was used for all experiments. Δ*fadR* and Δ*fadF* strains were constructed using lambda-red recombination (40). *fabB* and *fabA* were amplified from *E. coli* genomic DNA and cloned into BglBrick plasmid pBbA2k (41) using restriction digestion and ligation. All primers used are listed in **Supplemental Table 1.**

### LCMS analysis

Culture sampling and analyses of acyl-ACP, phospholipids, and proteins were performed largely as described in reference (17). In brief, samples were removed from cultures and quenched by adding directly to a 10% solution of trichloroacetic acid (2% final concentration). Quenched samples were pelleted by centrifugation and stored at -80 °C until analysis. For acyl-ACP and proteomics analysis, samples were lysed by suspending quenched pellets in a lysis solution with ^15^N-labelled internal standards before protein precipitation and digestion by Glu-C protease (acyl-ACP analysis) or trypsin (proteomics analysis). For phospholipid analysis, phospholipids were extracted in MTBE solution (42, 43) pre-mixed with internal standards consisting of a phospholipid extract from a ^13^C- labelled culture. LC/MS quantification followed the methods described in (17).

### Microplate experiments

All microplate experiments were performed with a Biotek Synergy HTX using 6-well plates. For experiments performed at stable temperatures, 2 mL of media was inoculated and plates continuously agitated (180 cm, 6 mm amplitude) with optical density (600 nm) recorded every 10 minutes. For cold shock experiments, exponential-phase cultures were prepared in stir flasks maintained at 37 °C before transfer to Falcon tubes on ice. After 10 minutes of incubation, 2 mL were transferred to microplate wells and growth monitored using the microplate reader. For cold shock experiments, plates were continuously agitated as before with optical density recorded every 5 minutes.

### Mathematical modelling

Steady-state and dynamic simulations were performed using a mathematical model constructed using Mathematica 13.2. Full details are provided in the **Supplemental Text**.

## Notes

### Competing Interest Statement

The authors have declared no competing interest.

## References

1. T. Harayama, H. Riezman, Understanding the diversity of membrane lipid composition. Nat. Rev. Mol. Cell Biol. 2018 195 **19**, 281–296 (2018).

2. M. Sinensky, Homeoviscous adaptation: a homeostatic process that regulates the viscosity of membrane lipids in Escherichia coli. Proc. Natl. Acad. Sci. U. S. A. 71 (1974).

3. J. B. Parsons, C. O. Rock, Bacterial lipids: Metabolism and membrane homeostasis. Prog. Lipid Res. 52, 249–276 (2013).

4. D. De Mendoza, Temperature sensing by membranes. Annu. Rev. Microbiol. 68, 101–116 (2014).

5. R. Ernst, S. Ballweg, I. Levental, Cellular mechanisms of physicochemical membrane homeostasis. Curr. Opin. Cell Biol. 53, 44–51 (2018).

6. P. E. Tiku, A. Y. Gracey, A. I. Macartney, R. J. Beynon, A. R. Cossins, Cold-induced expression of delta 9-desaturase in carp by transcriptional and posttranslational mechanisms. Science 271, 815–818 (1996).

7. P. S. Aguilar, A. M. Hernandez-Arriaga, L. E. Cybulski, A. C. Erazo, D. De Mendoza, Molecular basis of thermosensing: a two-component signal transduction thermometer in Bacillus subtilis. EMBO J. 20, 1681–1691 (2001).

8. S. Ballweg, et al., Regulation of lipid saturation without sensing membrane fluidity. Nat. Commun. 2020 111 **11**, 1–13 (2020).

9. P. S. Aguilar, J. E. Cronan, D. De Mendoza, A Bacillus subtilis gene induced by cold shock encodes a membrane phospholipid desaturase. J. Bacteriol. 180, 2194–2200 (1998).

10. S. Ballweg, R. Ernst, Control of membrane fluidity: The OLE pathway in focus. Biol. Chem. 398, 215–228 (2017).

11. J. E. Cronan, Thermal regulation of the membrane lipid composition of Escherichia coli. Evidence for the direct control of fatty acid synthesis. J. Biol. Chem. 250, 7074–7077 (1975).

12. K. Zhu, Y. M. Zhang, C. O. Rock, Transcriptional Regulation of Membrane Lipid Homeostasis in Escherichia coli. J. Biol. Chem. 284, 34880–34888 (2009).

13. J. L. Garwin, J. E. Cronan, Thermal modulation of fatty acid synthesis in Escherichia coli does not involve de novo enzyme synthesis. J. Bacteriol. 141, 1457–1459 (1980).

14. J. L. Garwin, A. L. Klages, J. E. Cronan, Structural, enzymatic, and genetic studies of β- ketoacyl-acyl carrier protein synthases I and II of Escherichia coli. J. Biol. Chem. 255, 11949– 11956 (1980).

15. A. K. Ulrich, D. de Mendoza, J. L. Garwin, J. E. Cronan Jr., Genetic and biochemical analyses of Escherichia coli mutants altered in the temperature-dependent regulation of membrane lipid composition. J. Bacteriol. 154, 221–230 (1983).

16. M. J. Noga, et al., Mass-Spectrometry-Based Quantification of Protein-Bound Fatty Acid Synthesis Intermediates from Escherichia coli. J. Proteome Res. 15, 3617–3623 (2016).

17. M. J. Noga, et al., Posttranslational Control of PlsB Is Sufficient To Coordinate Membrane Synthesis with Growth in Escherichia coli. MBio 11, e02703–19 (2020).

18. S. E. Goelz, J. E. Cronan Jr., The positional distribution of fatty acids in Escherichia coli phospholipids is not regulated by sn-glycerol 3-phosphate levels. J. Bacteriol. 144, 462–464 (1980).

19. Y. Xu, R. J. Heath, Z. Li, C. O. Rock, S. W. White, The FadR.DNA complex. Transcriptional control of fatty acid metabolism in Escherichia coli. J. Biol. Chem. 276, 17373–17379 (2001).

20. X. Xiao, X. Yu, C. Khosla, Metabolic flux between unsaturated and saturated fatty acids is controlled by the FabA:FabB ratio in the fully reconstituted fatty acid biosynthetic pathway of Escherichia coli. Biochemistry 52, 8304–8312 (2013).

21. I. Budin, et al., Viscous control of cellular respiration by membrane lipid composition. Science *(80-.).* **362** (2018).

22. K. Mains, J. Peoples, J. M. Fox, Kinetically guided, ratiometric tuning of fatty acid biosynthesis. Metab. Eng. 69, 209 (2022).

23. P. Edwards, J. S. Nelsen, J. G. Metz, K. Dehesh, Cloning of the fabF gene in an expression vector and in vitro characterization of recombinant fabF and fabB encoded enzymes from Escherichia coli. FEBS Lett. 402, 62–66 (1997).

24. E. P. Gelmann, J. E. Cronan, Mutant of Escherichia coli deficient in the synthesis of cis-vaccenic acid. J. Bacteriol. 112 (1972).

25. R. J. Heath, et al., Mechanism of triclosan inhibition of bacterial fatty acid synthesis. J. Biol. Chem. 274, 11110–11114 (1999).

26. Y. M. Zhang, H. Marrakchi, C. O. Rock, The FabR (YijC) transcription factor regulates unsaturated fatty acid biosynthesis in Escherichia coli. J. Biol. Chem. 277, 15558–15565 (2002).

27. M. B. Jackson, J. E. Cronan, An estimate of the minimum amount of fluid lipid required for the growth of escherichia coli. Biochim. Biophys. Acta - Biomembr. 512, 472–479 (1978).

28. C. Lau, et al., Conditions influencing formation of 16:0/16:0 molecular species in membrane phospholipids of Escherichia coli. J. Biol. Chem. 258 (1983).

29. W. Klein, M. H. W. Weber, M. A. Marahiel, Cold shock response of Bacillus subtilis: isoleucine-dependent switch in the fatty acid branching pattern for membrane adaptation to low temperatures. J. Bacteriol. 181, 5341–5349 (1999).

30. M. K. Shaw, J. L. Ingraham, Fatty Acid Composition of Escherichia coli as a Possible Controlling Factor of the Minimal Growth Temperature . J. Bacteriol. 90 (1965).

31. Y. Feng, J. E. Cronan, Complex binding of the FabR repressor of bacterial unsaturated fatty acid biosynthesis to its cognate promoters. Mol. Microbiol. 80, 195–218 (2011).

32. U. Alon, An introduction to systems biology: Design principles of biological circuits (2006).

33. D. K. Fujii, A. J. Fulco, Biosynthesis of unsaturated fatty acids by bacilli. Hyperinduction and modulation of desaturase synthesis. J. Biol. Chem. 252, 3660–3670 (1977).

34. F. J. Lombardi, A. J. Fulco, Temperature-mediated hyperinduction of fatty acid desaturation in pre-existing and newly formed fatty acids synthesized endogenously in Bacillus megaterium. Biochim. Biophys. Acta 618, 359–363 (1980).

35. R. Grau, D. de Mendoza, Regulation of the synthesis of unsaturated fatty acids by growth temperature in Bacillus subtilis. Mol. Microbiol. 8, 535–542 (1993).

36. D. S. Nichols, K. A. Presser, J. Olley, T. Ross, T. A. McMeekin, Variation of branched-chain fatty acids marks the normal physiological range for growth in Listeria monocytogenes. Appl. Environ. Microbiol. 68, 2809–2813 (2002).

37. A. K. Singh, et al., FabH selectivity for anteiso branched-chain fatty acid precursors in low-temperature adaptation in Listeria monocytogenes. FEMS Microbiol. Lett. 301, 188–192 (2009).

38. L. P. Saunders, S. Sen, B. J. Wilkinson, C. Gatto, Insights into the mechanism of homeoviscous adaptation to low temperature in branched-chain fatty acid-containing bacteria through modeling fabh kinetics from the foodborne pathogen listeria monocytogenes. Front. Microbiol. 7, 13861 (2016).

39. F. C. Neidhardt, P. L. Bloch, D. F. Smith, Culture Medium for Enterobacteria. J. Bacteriol. 119, 736–747 (1974).

40. K. A. Datsenko, B. L. Wanner, One-step inactivation of chromosomal genes in Escherichia coli K-12 using PCR products. Proc. Natl. Acad. Sci. U. S. A. 97, 6640–6645 (2000).

41. T. Lee, et al., BglBrick vectors and datasheets: A synthetic biology platform for gene expression. J. Biol. Eng. 5, 12 (2011).

42. V. Matyash, G. Liebisch, T. V Kurzchalia, A. Shevchenko, D. Schwudke, Lipid extraction by methyl-tert-butyl ether for high-throughput lipidomics. J. Lipid Res. 49, 1137–1146 (2008).

43. O. Susanto, et al., LPP3 mediates self-generation of chemotactic LPA gradients by melanoma cells. J. Cell Sci. 130, 3455–3466 (2017).

