## Supplemental Material for "A temperature-sensitive metabolic valve and a transcriptional feedback loop drive rapid homeoviscous adaptation in *Escherichia coli*"

### Supplemental Figure 1

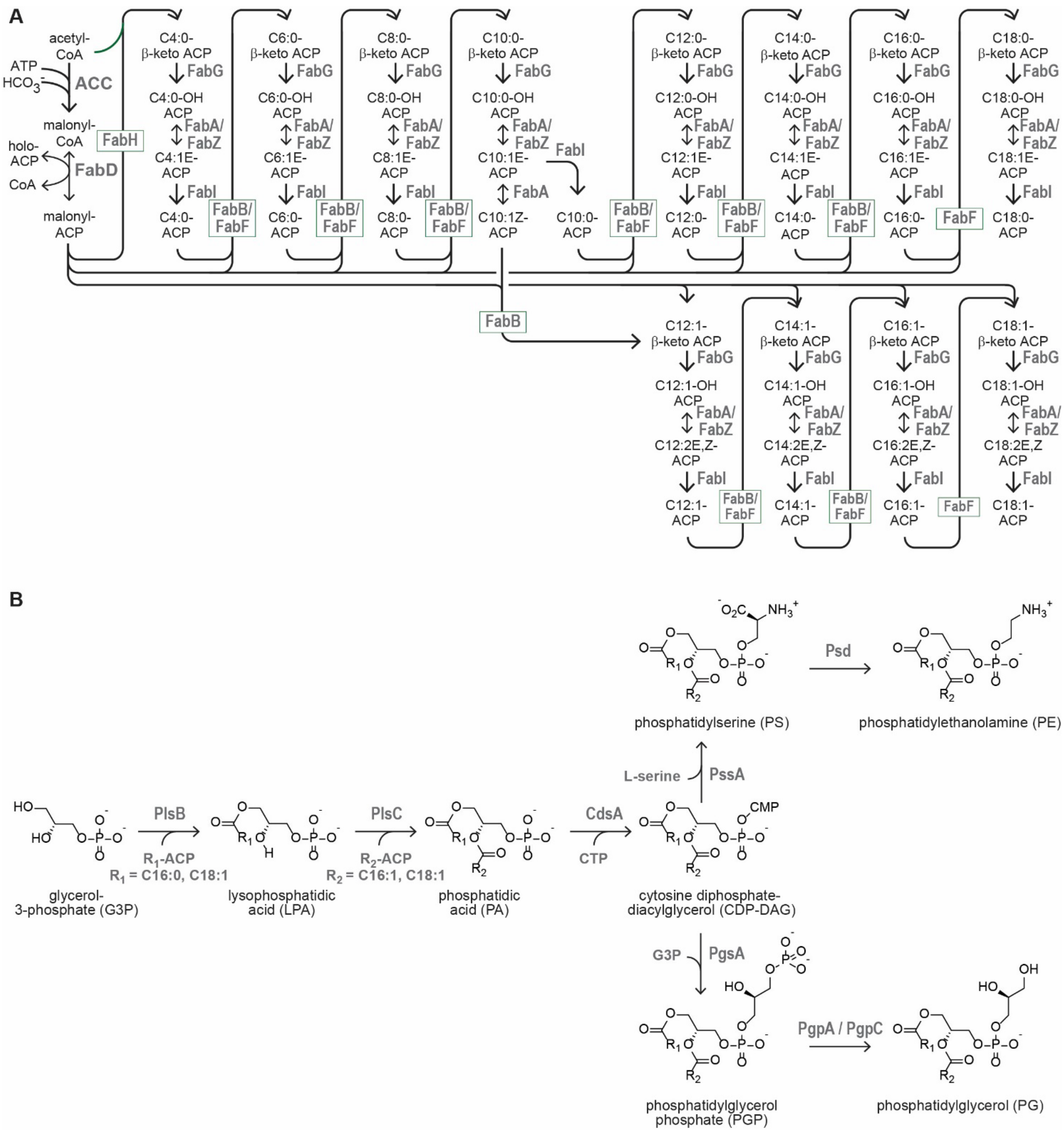


**Supplemental Figure 1.** The complete *E. coli* fatty acid **(A)** and phospholipid synthesis pathways **(B)**, indicating enzymes catalysing each reaction.

#### Supplemental Figure 2

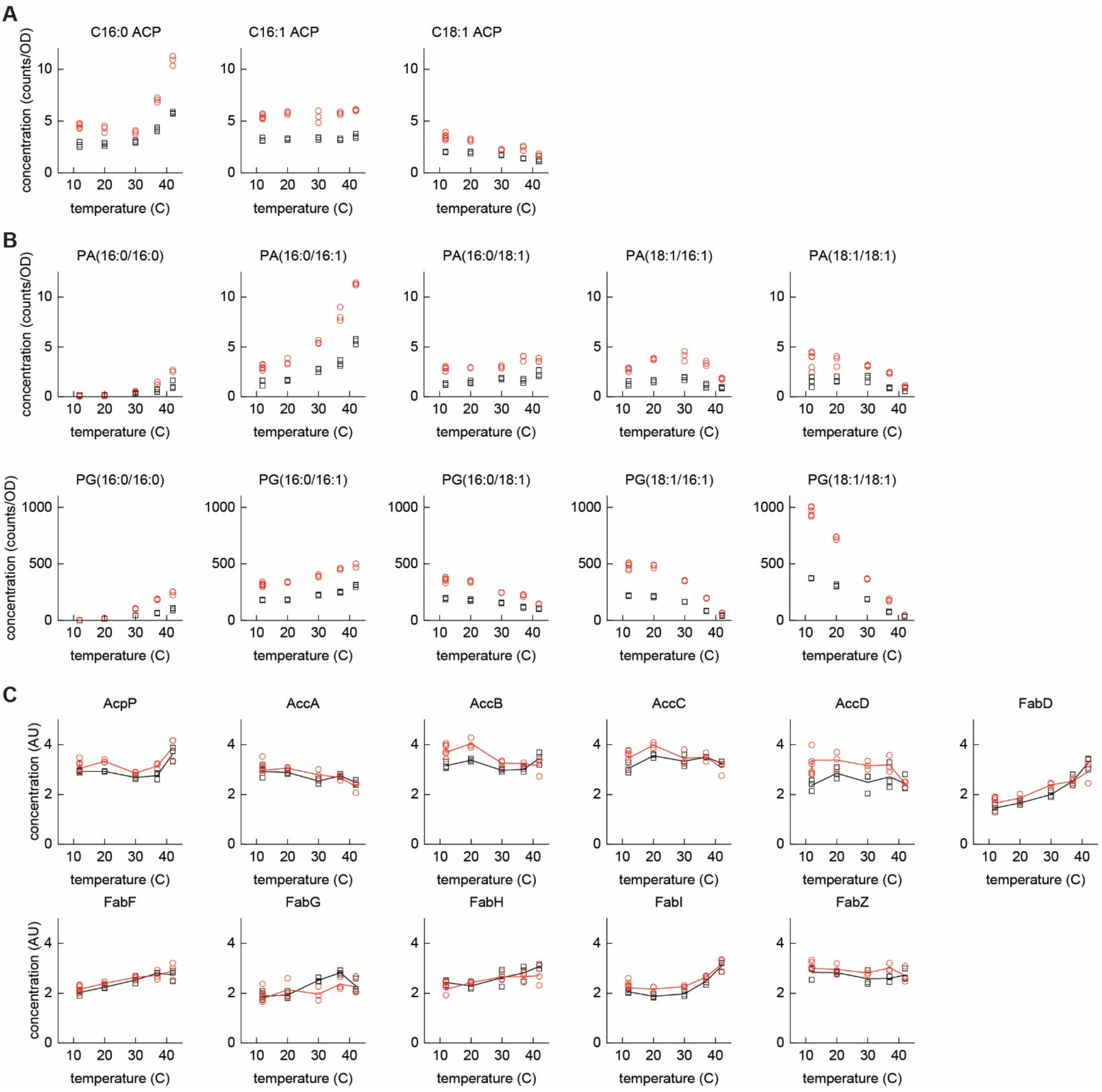


**Supplemental Figure 2.** Abundance of acyl-ACP **(A)**, phospholipid intermediates and membrane phospholipids **(B),** and enzymes of the fatty acid synthesis pathway **(C)** across 5 temperatures. Data from 3 measurements from two independent cultures at each temperature (black and red symbols).

#### Supplemental Figure 3

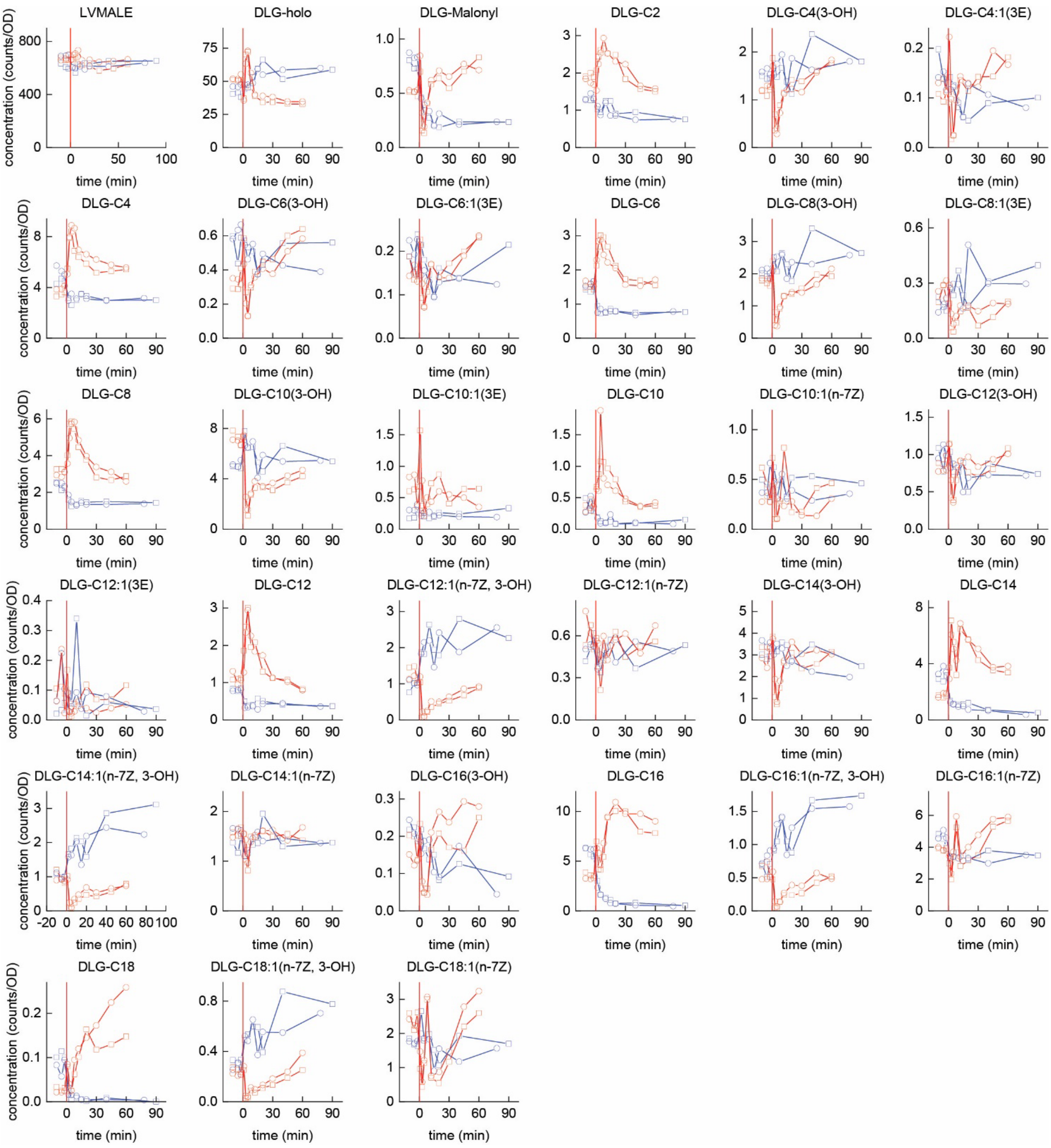


**Supplemental Figure 3.** Acyl-ACP concentrations following cold shocks (blue lines/symbols) and heat shocks (red lines/symbols). Two independent cultures are shown for each temperature shock (one measurement per time point).

#### Supplemental Figure 4

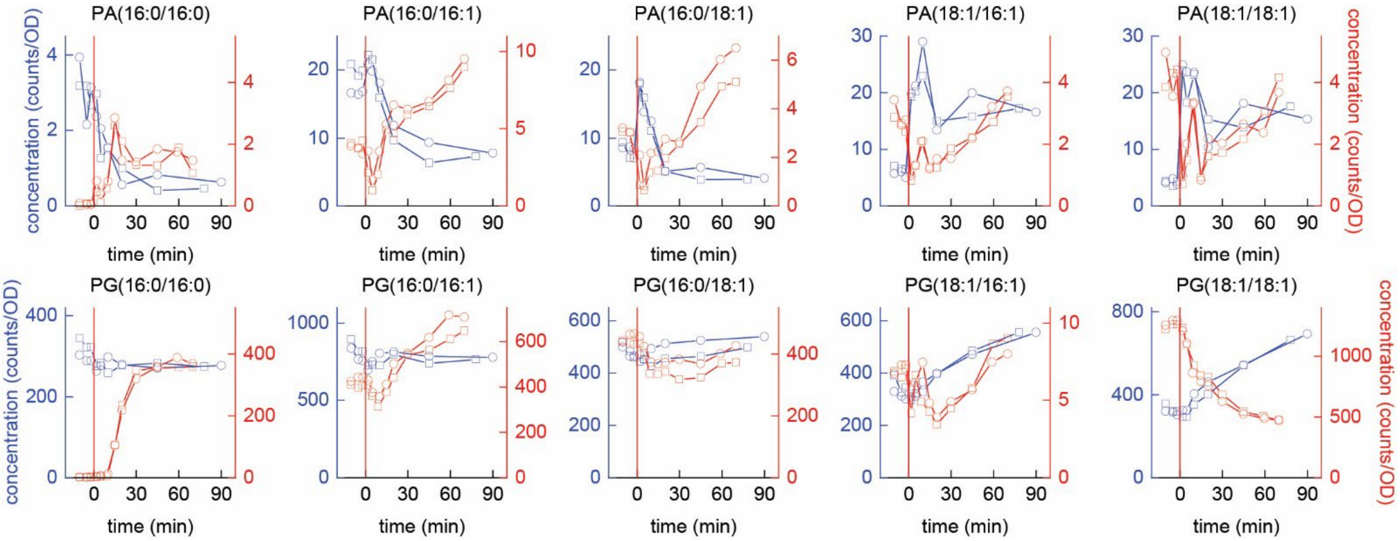


**Supplemental Figure 4.** Concentrations of phospholipid synthesis intermediates and membrane phospholipids following cold shocks (blue lines/symbols) and heat shocks (red lines/symbols). Data from 2 independent cultures are shown for each temperature shock.

#### Supplemental Figure 5

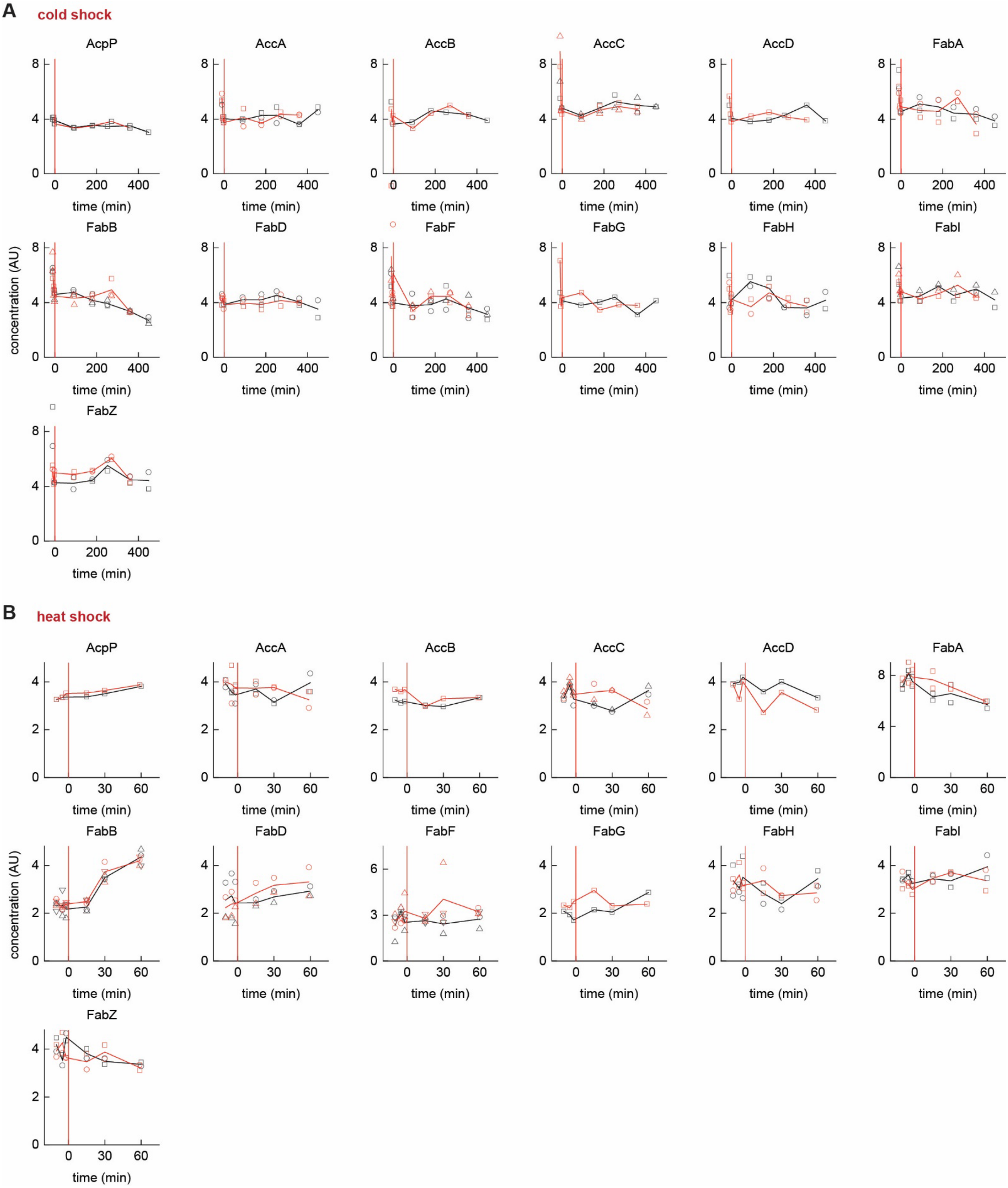


**Supplemental Figure 5.** Concentrations of fatty acid synthesis pathway enzymes following cold shocks **(A)** and heat shocks **(B).** Data for 2 independent cultures are shown for each temperature shock. Scatter points depict concentrations of individual peptides relative to an internal standard, while lines indicates averages of the peptide abundances for each replicate.

#### Supplemental Figure 6

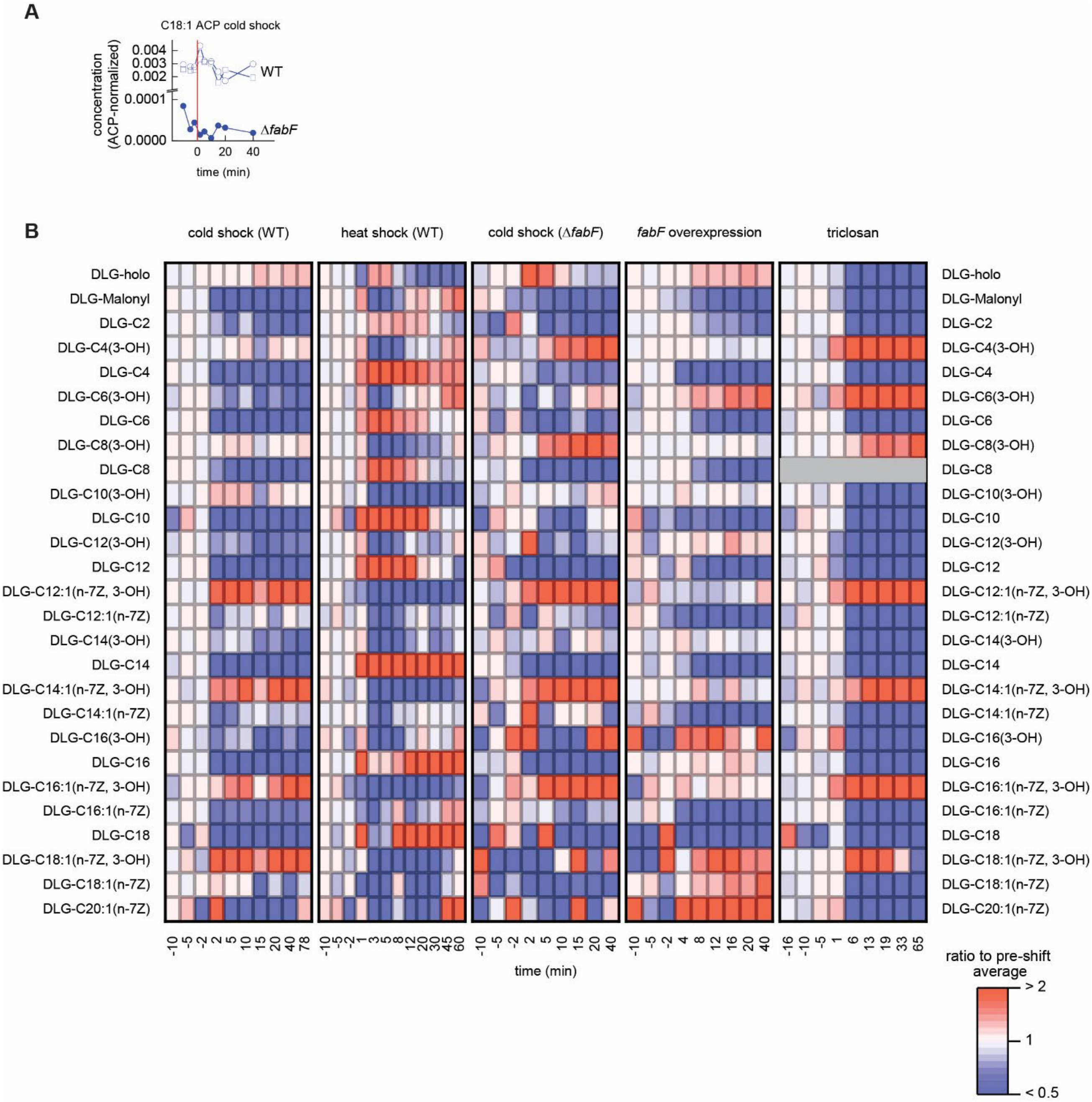


**Supplemental Figure 6. A.** C18:1 ACP quantification in *E. coli* Δ*fabF* and wild-type, indicating depleted abundance of C18:1 ACP in *E. coli* Δ*fabF*. **B.** Responses of acyl-ACP pools to cold shock (wild-type and Δ*fabF*), heat shock, *fabF* overexpression, and triclosan treatment. Each pool is normalized to the average value of the first three time points (pre-treatment). To facilitate comparison, one replicate is shown for each condition.

#### Supplemental Figure 7

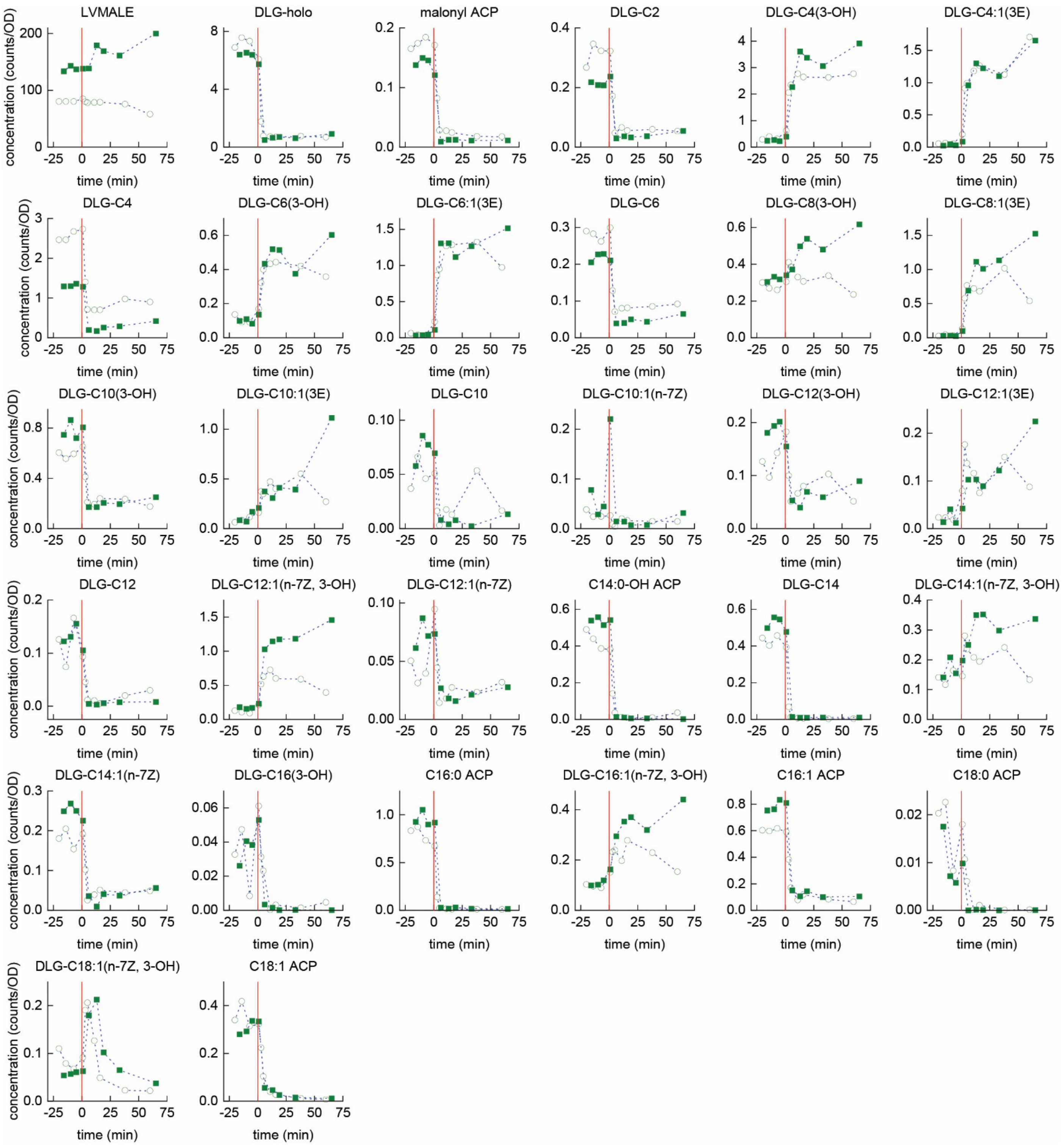


**Supplemental Figure 7.** Responses of acyl-ACP pools to triclosan. Data from 2 independent experiments are depicted.

#### Supplemental Text: Model description

We modelled the temperature response of the biosynthesis of saturated fatty acids (by FabI) and unsaturated fatty acids (by FabB) and their corresponding saturated and unsaturated phospholipid products with a minimal model. In addition to a temperature dependence of the catalytic rate constants of both FabI and FabB, this model considers the transcriptional regulation of FabB by its transcription factor FabR, whose activity depends on the concentration of saturated and and unsaturated fatty acids. We considered the dimerization of this transcription factor as well as the binding of its two regulatory fatty acids.

The model consists of the following differential equations:

$$\frac{ds(t)}{dt} = k_{FabI}(T)FABI - k_S s(t) \quad (1)$$

$$\frac{du(t)}{dt} = k_{FabB}(T)FabB(t) - k_U u(t) \quad (2)$$

$$\frac{dps(t)}{dt} = k_S s(t) - \mu ps(t) \quad (3)$$

$$\frac{dpu(t)}{dt} = k_U u(t) - \mu pu(t) \quad (4)$$

$$\frac{dFabB(t)}{dt} = \frac{k_t}{1+\left(\frac{FabR_2U_2(t)}{K_r}\right)^2} - \mu\, FabB(t) \quad (5)$$

$$FabR_2U_2(t) = \frac{\left(2FabR_T+K_{FabR}{}^2-\sqrt{K_{FabR}{}^2\left(4FabR_T+K_{FabR}{}^2\right)}\right)K_S{}^2u(t)^2}{2(K_SK_U+K_Us(t)+K_Su(t))^2} \quad (6)$$

$$k_i(T;\alpha_i,\beta_i,\gamma_i) = \frac{e^{-\frac{(T-\alpha_i)^2}{2\beta_i{}^2}}\ Erfc\left[-\frac{\gamma_i(T-\alpha_i)}{\beta_i\sqrt{2}}\right]}{\beta_i\sqrt{2\pi}}$$ (Erfc[x] is the complementary error function) (7)

Supplemental Figure 8 shows the temperature dependency of the catalytic rate constants $k_{FabI}(T)$ and $k_{FabB}(T)$ as function of temperature, using $\alpha_{FabI} = 320, \beta_{FabI} = 20, \gamma_{FabI} = -30, \alpha_{FabB} = 275, \beta_{FabB} = 15, \gamma_{FabI} = 30$.

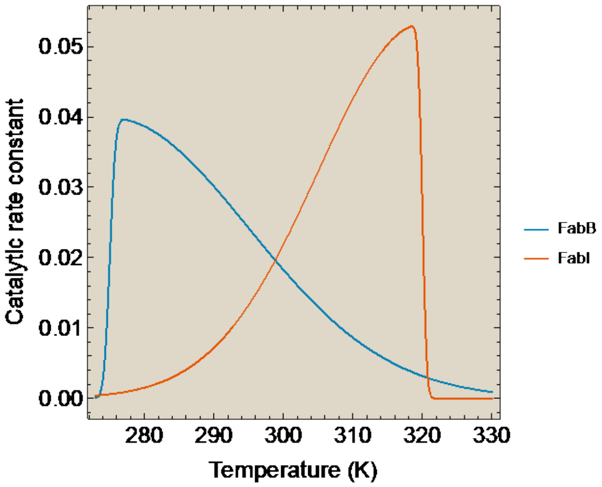


Supplemental Figure 8. The catalytic constants of FabI and FabB as function of temperature.

The model that describes the bound states of the transcription factor FabR with unsaturated and saturated fatty acid as function of the concentrations of those fatty acids, described both the dimerization of the transcription factor and the fatty acid binding events.

The dimerization is described by

$$FabR_T = FabR + 2\ FabR_{2T} \quad (8)$$

$$FabR_{2T} = \frac{FabR^2}{K_{FABR}} \quad (9)$$

which leads to the following equilibrium concentration of the dimer,

$$FabR_{2T} = \frac{1}{8}\left(4\ FabR_T + K_{FABR} - \sqrt{K_{FABR}(8\ FabR_T + K_{FABR})}\right) \quad (10)$$

This total dimer concentration occurs in different bound states,

$$FabR_{2T} = FabR_2 + 2FabR_2S + 2FabR_2U + 2FabR_2US + FabR_2S_2 + FabR_2U_2 \quad (11)$$

And the following associated binding equilibrium relations

$$FabR_2S = FabR_2\frac{s(t)}{K_S}, \quad (12)$$

$$FabR_2S = FabR_2\frac{u(t)}{K_S}, \tag{13}$$

$$FabR_2US = FabR_2\frac{u(t)s(t)}{K_SK_U} \tag{15}$$

$$FabR_2U_2 = FabR_2\left(\frac{u(t)}{K_U}\right)^2, \tag{16}$$

$$FabR_2S_2 = FabR_2\left(\frac{s(t)}{K_S}\right)^2 \tag{17}$$

All these equations can be solved for the transcription regulating state of FabR.

$$FabR_2U_2 = FabR_{2T}\frac{u(t)^2{K_S}^2}{\left(K_SK_U+K_Us(t)+K_Su(t)\right)^2} \tag{18}$$

This ends the description of the used model. Note that: 1) we do not know all the parameter values of this model and 2) the model captures the essence of the regulatory system and is therefore simplified and includes lumped processes and phenomenological parameters. We therefore simulated the model with normalized time and concentration units. One way to consider this is by writing the model in terms of dimensionless parameters and variables.

$$\frac{d\bar{s}(\tau)}{d\tau} = \kappa_{FabI}(T)\overline{FABI} - \kappa_S\bar{s}(\tau) \tag{19}$$

$$\frac{d\bar{u}(\tau)}{d\tau} = \kappa_{FabB}(T)\overline{FabB}(t) - \kappa_U\bar{u}(\tau) \tag{20}$$

$$\frac{d\overline{ps}(\tau)}{d\tau} = \kappa_S\bar{s}(\tau) - \overline{ps}(\tau) \tag{21}$$

$$\frac{d\overline{pu}(\tau)}{d\tau} = \kappa_U\bar{u}(\tau) - \overline{pu}(\tau) \tag{22}$$

$$\frac{d\overline{FabB}(\tau)}{d\tau} = \frac{\kappa'_t}{1+(\overline{FabR_2U_2}(\tau))^2} - \overline{FabB}(\tau) \tag{23}$$

$$\overline{FabR_2U_2}(\tau) = \alpha\frac{\left(4\,\beta+1-\sqrt{8\,\beta+1}\right)\bar{u}(\tau)^2}{8(1+\bar{s}(\tau)+\bar{u}(\tau))^2} \tag{24}$$

Dimensionless parameter and variable definitions:

$\tau = \mu t, \kappa_{FabI}(T) = k_{FabI}(T)/\ \mu, \kappa_S = k_S/\mu, \bar{s}(\tau) = s(\tau)/K_S, \overline{FABI} = FABI/K_S, \bar{u}(\tau) = u(\tau)/K_U,$

$\overline{FabB(t)} = FabB(t)/K_U, \kappa_U = k_U/\mu, \overline{ps}(\tau) = ps(\tau)/K_S, \overline{pu}(\tau) = pu(\tau)/K_U,$

$\kappa'_t = k_t/(\mu K_U), \overline{FabR_2U_2}(\tau) = FabR_2U_2(t)/K_r, \alpha = \frac{K_{FABR}}{K_r}, \beta = \frac{FabR_T}{K_{FABR}}.$

The used kinetic parameter values are: $k_s = 0.1\ (time^{-1}), k_U = 0.1\ (time^{-1}), \mu = 0.001(time^{-1}), k_t = 0.0005\ (conc\ time^{-1}), FabI = 0.5\ (conc), K_{FABR} = 0.1\ (conc), FabR_T = 1\ (conc), K_S = 0.1\ (conc), K_U = 0.05\ (conc), K_r = 0.1(conc)$ and therefore: $\kappa_S = 100, \kappa_U = 100, FabI = 5$, $\kappa'_t = 100$, $\alpha = 1$, $\beta = 10$. Thus, in the end the model has these 6 dimensionless parameters plus the 6 parameters for the temperature dependency of the catalytic rate constants FabI and FabB.

#### Supplemental Table 1

| Supplemental Table 1. Primer sequences | | |
|---|---|---|
| Name | Sequence | Used for |
| GB220426-fabF.f | aaaGAATTC GGAGGACAAACgtgTCTAAGCG | fabF cloning |
| GB220426-fabF.r | ttactcgagTTAGGATCCttaGATCTTTTTAAAGATCAAAGAACCATTAGTGC | fabF cloning |
| GB220426-fabB.f | aaaAGATCT GCGACTTACAGAGGTATTGAatgAAAC | fabB cloning |
| GB220426-fabB.r | ttactcgagTTAGGATCCttaATCTTTCAGCTTGCGCATTACC | fabB cloning |
| GB220426-fabA.f | aaaAGATCT AAATAAGGCTTACAGAGAACatgGTAG | fabA cloning |
| GB220426-fabA.r | ttactcgagTTAGGATCCtcaGAAGGCAGACGTATCCTGG | fabA cloning |
| GB230102fabFKO.f1 | CTAGAATCATTTTTTCCCTCCCTGGAGGACAAACgtg gtgtaggctggagctgcttc | fabF knockout |
| GB230102fabFKO.f2 | GTCGTTCGACCGCCTGAGTTTTATCTTTTTGTCCCACTAGAATCATTTTTTCCCTCCC | fabF knockout |
| GB230102fabFKO.r1 | GTGGAAAATGACAACttaGATCTTTTTAAAGATCAAatgggaattagccatggtcc | fabF knockout |
| GB230102fabFKO.r2 | GGCCCGCAAGCGGACCTTTTATAAGGGTGGAAAATGACAACttaGATC | fabF knockout |
| GB230105-fabFKO.fseq | CAGGCGTAAGTGAACATCTCC | confirming fabF knockout |
| GB230105-fabFKO.rseq | GTTCACTACGGAACAAGTCGG | confirming fabF knockout |
| GB230102fabRKO.f1 | CCGTTCATTCACAATACTGGAGCAATCCAGTatg gtgtaggctggagctgcttc | fabR knockout |
| GB230102fabRKO.f2 | CAAAGAATTGCAGTAAATATGTTTTATTGCGTTACCGTTCATTCACAATACTGG | fabR knockout |
| GB230102fabRKO.r1 | GCTTGTTTCAttaCTCGTCCTTCACATTTCCatgggaattagccatggtcc | fabR knockout |
| GB230102fabRKO.r2 | CTAACGCCAGCAGCAGCGTACCTCTATCTTGATTTGCTTGTTTCAttaCTCGTCC | fabR knockout |
| GB230105-fabRKO.fseq | GCAGGTTTCCGCTGTACG | confirming fabR knockout |
| GB230105-fabRKO.rseq | AGTACCATTAATCGATAAGCCAGC | confirming fabR knockout |
